# Analyzing the miRNA regulatory landscape of OGT identifies bidirectional regulation

**DOI:** 10.1101/2025.08.21.671361

**Authors:** Zainab Ali Syeda, Faezeh Jame-Chenarboo, Hoi-Hei Ng, Chu Thu, Ramiz Demir, Helia Dehghan Harati, Dawn Macdonald, Lara K. Mahal

## Abstract

O-GlcNAc transferase (OGT) is the key enzyme involved in post-translationally modifying cytoplasmic and nuclear proteins with O-GlcNAc. Maintenance of cellular O-GlcNAcylation levels is critical to cell health and requires precise transcriptional and post-transcriptional control. Herein we examine the miRNA regulation of OGT by the human miRNAome using our high-throughput miRFluR assay. We found >200 miRNA regulators of OGT, including 17 down- and 15 upregulatory miRNAs previously identified in CLIP datasets. We validated the impact of select miRNA on OGT and O-GlcNAc levels using both miRNA mimics and inhibitors that reduce endogenous miRNA levels. We focused our studies on two miRNA families, the downregulatory let-7 family and the upregulatory miR-148/152 family. For the let-7 family, we found that only let-7a-2-3p and let-7g-3p strongly downregulated OGT. Downregulation required two seed-dependent binding sites. Evolutionary analysis found that the more recent of the two sites emerged in placental mammals. All three members of the miR-148/152 family upregulated OGT. Although two putative sites for the family were identified in CLIP datasets, our results indicate that these sites alone are insufficient for OGT upregulation by miR-148/152. Rather, OGT is upregulated via multiple binding sites distributed throughout its 3’ UTR. Overall, our results provide important insights into miRNA mediated regulation of OGT,and reveal multi-site regulation by both upregulatory and downregulatory miRNAs.

## Introduction

O-GlcNAc, in which *N-*acetylglucosamine modifies serines or threonines, is a nucleocytoplasmic modification that is crucial for cell cycle signaling, stress response and metabolic sensing (1). Levels of O-GlcNAc are controlled by O-GlcNAc transferase (OGT) (2, 3), which puts on the modification, and O-GlcNAcase (OGA)(4–10), which removes it (**Fig. 1**)(10–13). Perturbations in O-GlcNAc have been observed in neurodegenerative disorders such as Alzheimer’s and Parkinson’s disease, Type 2 diabetes and cardiovascular disorders (14–16). This dynamic modification requires tight regulation of both the enzymes OGT and OGA, and UDP-GlcNAc levels.

**Figure 1.**
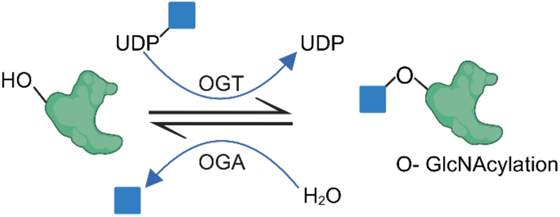
Scheme for O-GlcNAcylation. O-GlcNAc transferase (OGT) transfers *N-*acetyl-D-glucosamine (GlcNAc, blue square) onto serines or threonines on proteins. The enzyme O-GlcNAcase (OGA) is responsible for the removal of this modification, making it reversible. UDP-GlcNAc: uridine diphosphate *N*-acetyl-D-glucosamine.

OGT is a highly conserved gene across most eukaryotes, from C. *elegans* to humans (17–19). Conservation of this enzyme suggests that O-GlcNAc plays an ancient and fundamental role. OGT is essential for embryogenesis and the viability of dividing cells (20, 21). In mammals, alternative splicing produces three *OGT* isoforms: nucleocytoplasmic *OGT* (ncOGT), mitochondrial *OGT* (mOGT), and short *OGT* (sOGT), which differ in location and transcript length but share the same 3’ untranslated region (UTR), the primary site of miRNA regulatory interactions (22).

miRNAs are endogenous non-coding RNAs (∼22 nucleotides in length) that fine tune protein expression. Canonically, they are thought to work by inhibiting translation or causing degradation of messenger RNA (mRNA) (23–25) through complementation of 6-7 nucleotides between mRNA and the seed-region of the miRNA. It has become increasingly clear that miRNA regulation is bidirectional, with the same miRNA inhibiting the expression of one protein, while enhancing that of another (26–29). First identified in non-dividing cells in 2007 by Vasudevan and Steitz, upregulation of protein expression by miRNA has been observed as a common occurrence in actively dividing cells (29–34). In contrast to downregulation, upregulation often utilizes non-canonical (non-seed) binding between the miRNA and mRNA (26–29). While the precise molecular mechanisms of upregulation are still unclear, data has shown a requirement for Argonaute 2 (AGO2), but not the trinucleotide repeating-containing adaptor 6A (TNRC6A, aka GW182), a critical part of all downregulatory AGO2 complexes, in miRNA mediated upregulation (29, 32, 33, 35). At present, while seven miRNA have been identified as downregulators of OGT (36–42), it is unknown whether miRNA can upregulate this critical enzyme.

Herein, we did a comprehensive analysis of miRNA regulation of OGT via its 3’UTR using our high-throughput assay, miRFluR (26–30). We identified ∼200 regulators of OGT, including 17 down- and 15 upregulatory miRNAs previously found in CLIP datasets, identifying a bidirectional role for miRNA in tuning OGT expression. We focused on two miRNA families, the let-7 family, members of which downregulated OGT, and the miR-148/152 family, which upregulated OGT. In both cases, we found multi-site regulation of OGT, with two binding sites identified for the let-7 family and distributed multi-site regulation for the miR148/152 family. Overall, our work sheds light on the post-transcriptional regulation of this crucial enzyme.

## Results

### High-throughput miRFluR analysis of OGT identifies both up- and downregulatory miRNA

O-GlcNAc transferase (OGT) is one of the most highly regulated proteins in the cell, with documented regulation at the transcriptional, post-transcriptional and post-translational levels (43–45). Levels of this enzyme are tuned by external environmental conditions, including stress, and are sensitive to metabolic perturbations which alter O-GlcNAc levels (46–48). Post-transcriptional regulation by miRNA has been observed and to date, a handful of miRNA regulators have been studied (37–42, 49, 50). To provide a more comprehensive view, we utilized our high-throughput miRFluR assay to test the human miRNAome for OGT regulators. This assay utilizes a genetically encoded ratiometric sensor (pFmiR) containing the fluorescent protein Cerulean upstream of the 3’-untranslated region (3’UTR) of the gene of interest and mCherry as an internal control (**Fig. 2A**). Co-transfection into HEK-293T cells of the plasmid and a library of miRNA allows interrogation of the miRNAome. Ratiometric analysis of Cerulean/mCherry fluorescence indicates the extent of miR:target regulation.

**Figure 2.**
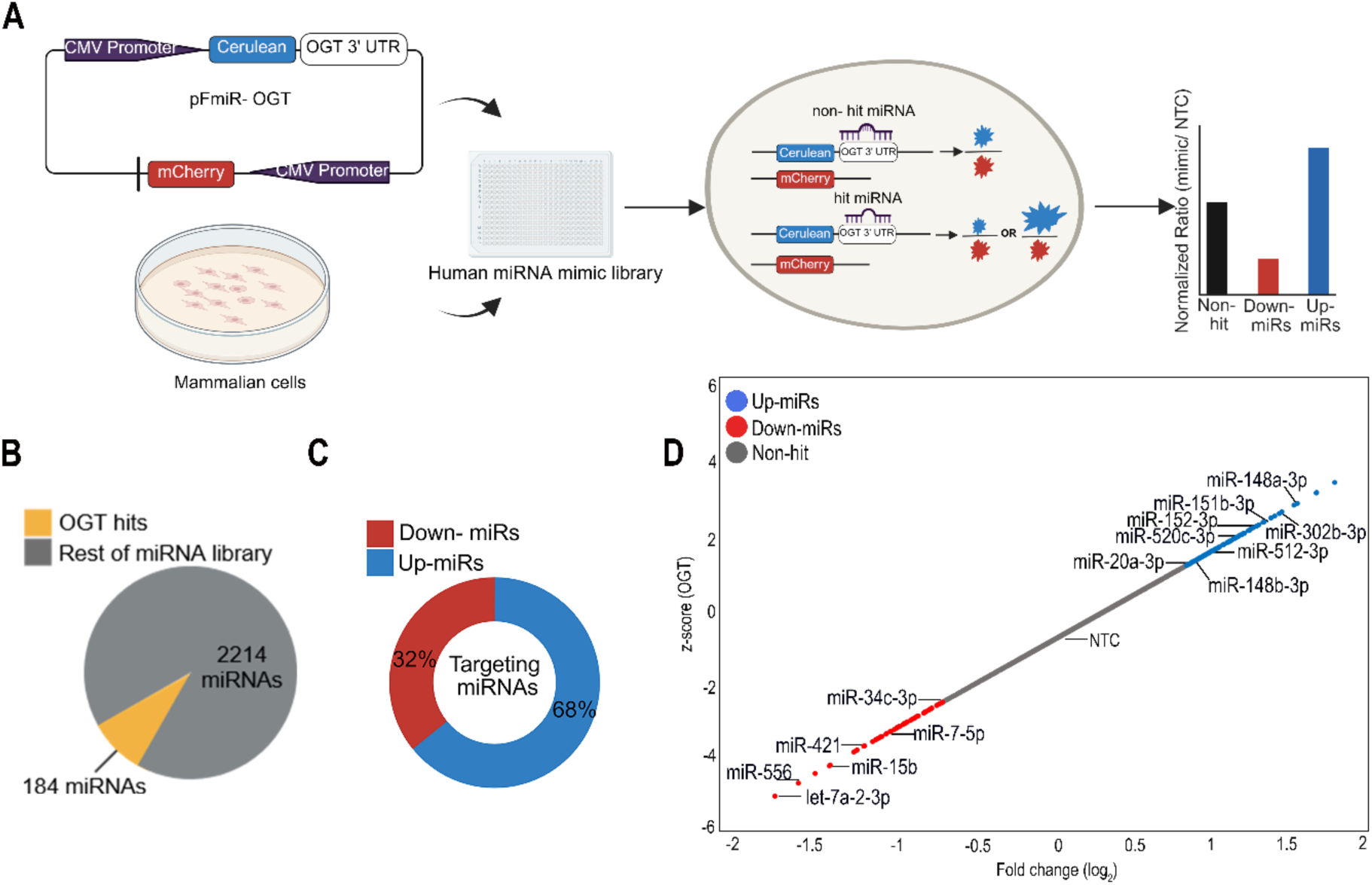
High throughput analysis of OGT 3’UTR using miRFluR assay. (A) Scheme of miRFluR assay. The pFmiR-OGT 3’UTR sensor is co-transfected with the human miRNA library in HEK -293T cells. The ratio of Cerulean to mCherry is normalized to non-targeting control (NTC) and can identify downregulatory (down-miR, red), neutral, or up-regulatory (up-miR, blue) miRNA. (B) Pie chart of miRNA hits (yellow, 8.3 %) for OGT 3’UTR. (C) Representation of up-miRs (blue, 68 %) versus down-miRs (red, 32 %) among miRNAs hits. (D) Scatter plot of Z-score versus fold-change (log2) for QCed miRFluR data. miRNAs in the 90 % confidence interval are colored (down-miRs: red, up-miRs: blue) and those selected for further validation are labeled. Associated data is shown in **Figs. S1-S2, Table S1**.

To study the miRNA regulation of OGT, we cloned the 3’UTR of human OGT into our pFmiR plasmid (**Figs.S1-S2**). Although OGT has multiple isoforms, all share a single conserved 3’UTR (2147 nucleotides (nt)). We co-transfected pFmiR-OGT plasmid with the human miRNA mimic library (Dharmacon, v. 21) into HEK-293T cells. The library was displayed in 384-well plates (24 total) with 3-replicates for each miRNA mimic. After 48 hours (h), we analyzed the fluorescence of the cells and calculated the Cerulean:mCherry ratio. We normalized the data to a non-targeting control (NTC) present in each plate. After quality control (QC), our dataset contained 2214 miRNAs (**Dataset S1, see Methods**). After z-scoring our data, we noted that several of the known downregulators of OGT were outside of the 95% confidence interval. Of the seven miRNA previously shown to downregulate OGT, one was QCed out (miR-424-5p) (36), one showed no change (miR-485-5p)(37) and five showed downregulation of at least 30% in our assay (miRs-7-5p, - 24-3p, -101-5p, -483-3p, -501-3p) (38–42). Only two of the five known hits were in the 95% confidence interval (miRs-7-5p, -501-3p). In general, literature using genetically encoded sensors uses 20% downregulation of signal (i.e. 0.8 miR/NTC ratio) as the cutoff for biological significance for miRNA regulation (30, 51). Taking into account the previous work, we decided to set the 90% confidence interval as the delineation for our hit list (downregulators: 0.69 miR/NTC, upregulators: 1.6 miR/NTC) as it encompassed more of the literature hits. We found that 8.3% of miRNA passing QC were in this interval (184 total hits, **Fig. 2B-D, Table S1**). This is significantly higher than what is observed in our other miRFluR data sets (3-6%) and is consistent with both our expanded confidence interval and the predictions that OGT is one of the most highly regulated genes by miRNA (26–29, 52). We found a higher number of upregulatory (**Fig. 2C**, up-miRs, 68%, 125 miRs) than downregulatory (down-miRs, 32%, 59 miRs) miRNAs targeting OGT. We compared our miRNA hit-list with data generated through cross-linked immunoprecipitation (CLIP) of AGO complexes using starBase v.2., a comprehensive CLIP database (53). In CLIP, mRNA footprinting is combined with miRNA profiling from the IP and miRNA prediction algorithms are used to determine binding sites (54). We found that 17 down-miRs and 15 up-miRs from our hit list were identified in AGO complexes with the OGT 3’UTR (**Datasets S1 & S2**), including 3 of the known down-miRs. This data confirms the ability of our miRFluR assay to identify miRNA regulation of OGT.

### miRNAs regulate OGT expression bidirectionally in cancer cell lines

To further validate our miRFluR results, we chose a set of miRNA (5 down-miRs, 6 up-miRs) from throughout the confidence interval and added the known miRNA regulator, miR-7-5p as a comparator (40, 55). We tested down-miRs: let-7a-2-3p, <u>miR-7-5p</u>, -15b, -556, -421, -34c-3p and up-miRs: <u>miR-302b-3p</u>, <u>-148a-3p</u>, -20a-3p, -151b-3p, - 512-3p, and <u>-520c-3p</u> (miRNAs observed in CLIP datasets are underlined, **Dataset S2**). We transfected mimics or NTC into A549, a common lung adenocarcinoma cell line, and analyzed OGT levels by Western blot analysis and RT-qPCR at 72 h post-transfection (**Fig. 3, Fig. S3, Table S2**). Of the six down-miRs tested, only two impacted OGT levels in this cell line when compared to NTC (let-7a-2-3p: 0.67-fold, miR-7-5p: 0.71-fold, **Fig. 3A-B, Fig. S3A-B**). In contrast, we observed clear upregulation for all of the up-miRs, although statistical significance was only reached for miR-148a-3p (miR-302b-3p: 1.8-fold, miR-148a-3p: 1.8 fold, miR-20a-3p: 2.1 fold, miR-151b-3p: 2.1 fold, miR-512-3p: 2.2 fold, miR-520c-3p: 1.8 fold). In line with our previous studies, the impact of miRNAs on the mRNA levels of OGT, as determined by RT-qPCR (**Fig. 3C, Table S2**), showed mixed results, with some changes concordant (let-7a-2-3p) and some discordant (miR-148a-3p) with the protein data.

**Figure 3.**
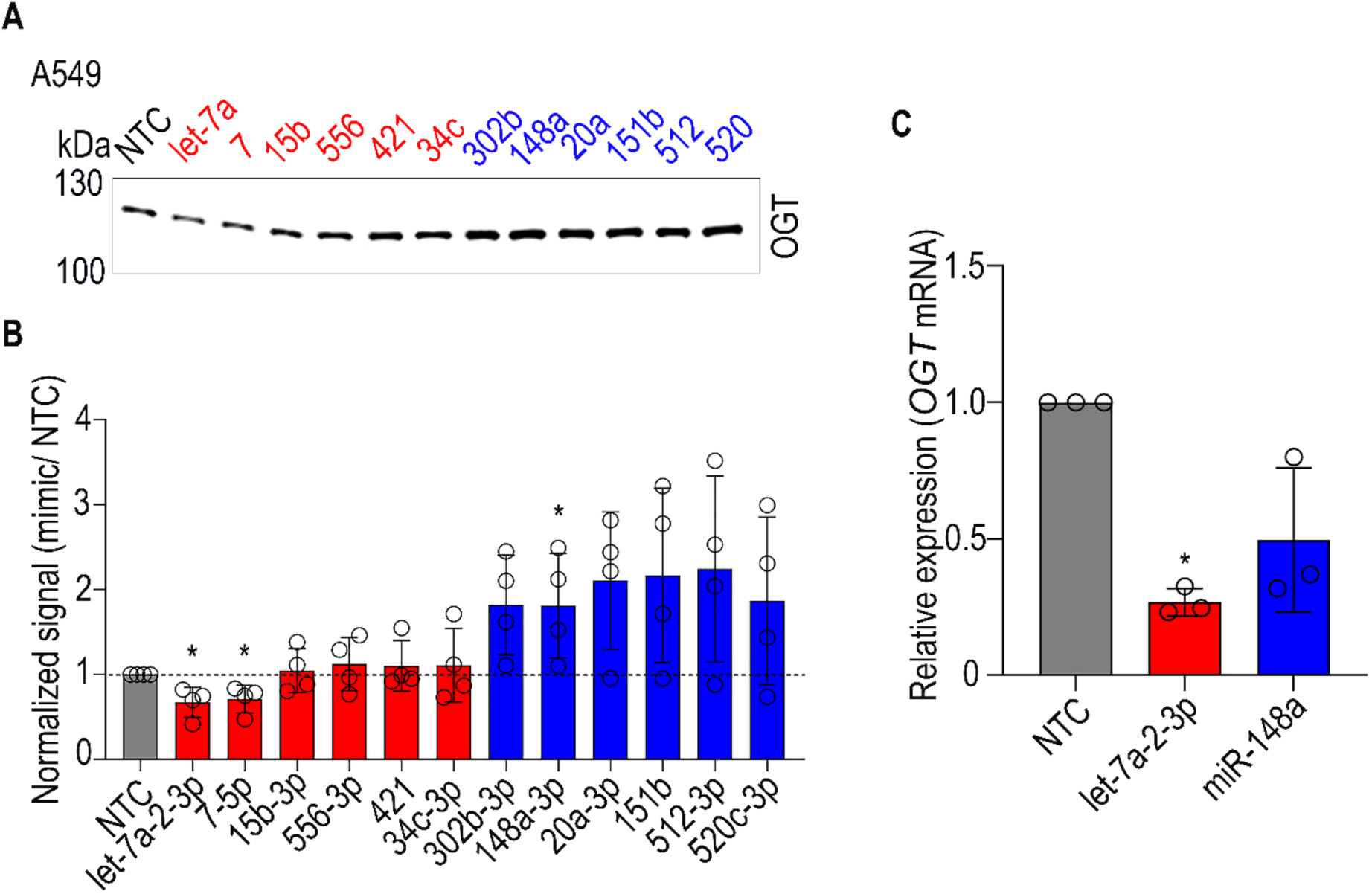
Validation of miRNA regulators of OGT in A549. (A) Representative Western blot analysis for α-OGT. A549 cells were treated with 50 nM of either non-targeting control (NTC) or miRNA mimics (down-miRs (red): -let-7a-2-3p, miR-7-5p, -15b-3p, -556-3p, - 421, -34c-3p. up-miRs (blue): miR-302b-3p, -148a-3p, -20a-3p, -151b-3p, -512-3p and - 520c-3p) for 72 h prior to lysis and analysis. Ponceau staining, the expanded blot and antibody validation are shown in **Fig. S3**. (B) Quantification of Western blot data for n=4 biological replicates. Western blot samples were normalized first to total protein (Ponceau) and then to the normalized NTC. Individual replicates are indicated as discrete points. Error bars represent standard deviations. (C) Quantification of RT-qPCR results for OGT in A549. A549 cells were transfected with miRNA mimics or NTC (50 nM, 72 h). Graph of normalized OGT mRNA expression data relative to GAPDH is shown (n=3 biological replicates). Primer sequence: **Table S3**. Statistical analysis was done using one-sample *t*-test (Western blot analysis) and paired *t*-test (Western blot analysis and RT-qPCR). Highest significance from either test is shown. *p*-values for both the tests are reported in **Table S2**. \**p < 0.05*.

We revalidated the impact of a subset of the miRNA on OGT levels in the colorectal adenocarcinoma line CaCo-2 and the neuroblastoma cell line SH-SY5Y (**Fig. 4, Fig. S4**; down-miRs: let-7a-2-3p*, miR-7-5p; up-miRs: miR-20a-3p, -148a-3p*, -302b-3p, *determined in both cell lines). In line with other studies, cell dependent effects were observed (50, 56–59). Let-7a-2-3p significantly downregulated OGT expression in both CaCo-2 (0.49 fold, **Fig. 4A-B**) and SH-SY5Y (0.60 fold, **Fig. 4C-D**). However, the known downregulator miR-7-5p had no impact on OGT in CaCo-2, arguing for cell line dependency in its action. In CaCo-2, only miR-20a-3p and -148a-3p were statistically significant upregulators of OGT (miR-20a-3p: 2.6 fold, miR-148a-3p: 3.0 fold). In contrast, in SH-SY5Y all of the biological replicates for miR-148a-3p upregulated protein expression, but it did not meet statistical significance due to the variability in the response (range:1.2-1.9 fold). Throughout our studies, we found that OGT levels were more variable in their response to miRNA mimics than other genes tested for both up- and downregulation (for an example see data for known downregulator miR-7-5p in **Figs. 3 and 5**). This may be due to the high sensitivity of OGT to various forms of stimuli, including stress and glucose levels (1, 46). We note, however, that our miRNA data was highly consistent in trend (e.g. upregulators enhanced protein expression, downregulators inhibited) arguing that these are true biological responses to miRNA regulation.

**Figure 4.**
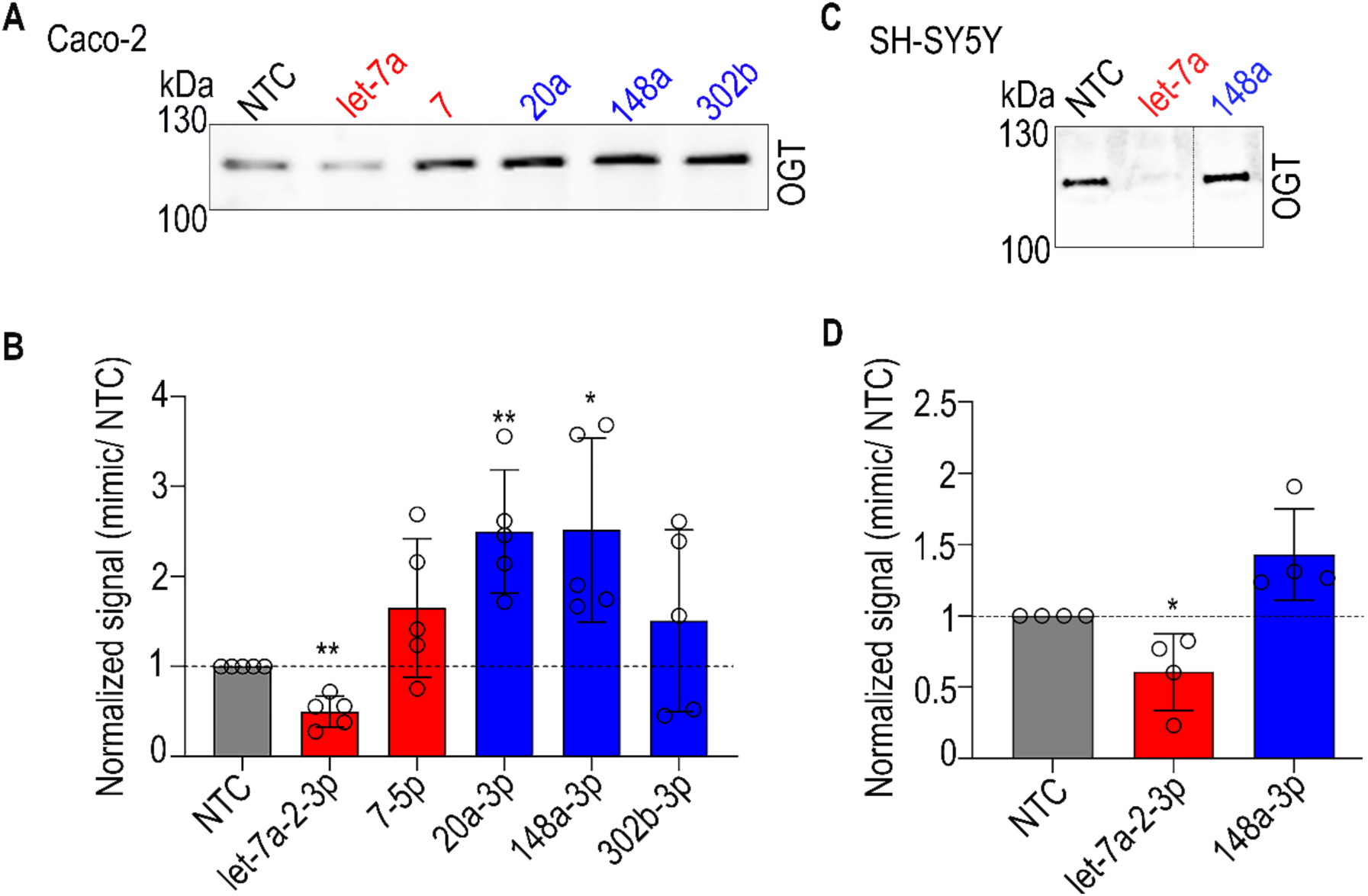
Validation of miRNA regulators of OGT in Caco-2 and SH-SY5Y. (A, C) Representative Western blot analysis for α-OGT. Caco-2 (A) and SH-SY5Y (C) cells were treated with 50 nM of either non-targeting control (NTC) or indicated miRNA mimics (down-miRs (red): let-7a-2-3p, miR-7-5p; up-miRs (blue): miR-20a-3p, -148a-3p and 302b-3p) for 72 h prior to lysis and analysis. (B, D) Quantification of Western blot data for Caco-2 (A) and SH-SY5Y (C) Bar charts represent average data for biological replicates (Caco-2: n=5, SH-SY5Y: n=4). Western blot samples were normalized first to total protein (Ponceau) and then to the normalized NTC. Individual replicates are indicated as discrete points. Error bars represent standard deviations. Additional associated data is shown in **Fig. S4**. Statistical analysis was done using both the one-sample *t*-test and paired *t*-test. Highest significance from either test is shown. *p*-values for both the tests are reported in **Table S2**. *\*p < 0.05, ** < 0.01*.

**Figure 5.**
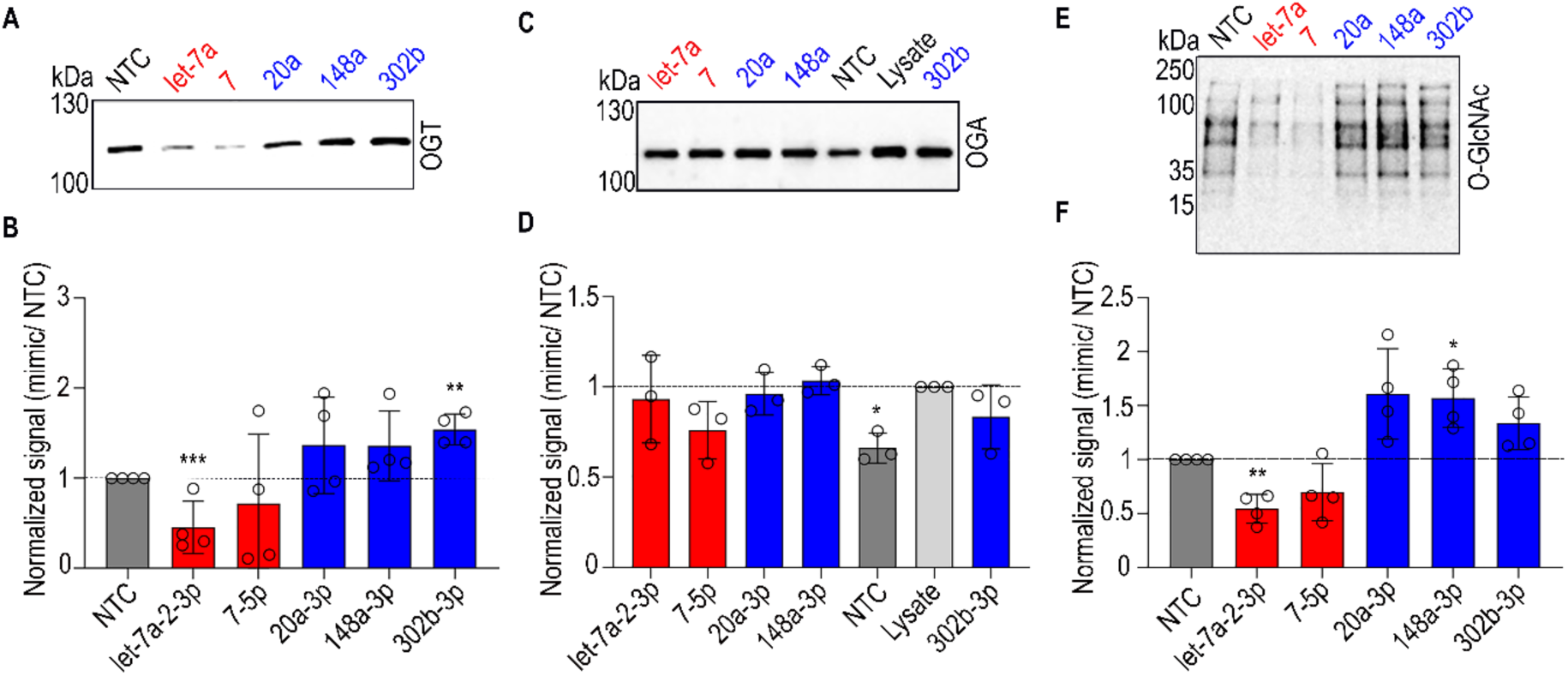
miRNAs regulate O-GlcNAc levels through OGT without impacting OGA. (A, C, E) Representative Western blot data for α-OGT (A), α-OGA (C) and α-O-GlcNAc (E). A549 cells were treated with 50 nM of either non-targeting control (NTC) or indicated miRNA mimics: NTC (grey), down-miRs (red): let-7a-2-3p, miR-7-5p, up-miRs (blue): miR-302b-3p, -148a-3p, -20a-3p; for 72 h prior to lysis and analysis. (B, D, F) Quantification of Western blot analysis for OGT (B, n=4), OGA (D, n=3) and O-GlcNAc (F, n=4). Bar charts represent average data for indicated biological replicates. For OGT and O-GlcNAc, Western blot samples were normalized first to total protein (Ponceau) and then to the normalized NTC. For OGA, untreated lysate was used for normalization in lieu of NTC. Individual replicates are indicated as discrete points. Error bars represent standard deviations. Additional associated data is shown in **Fig. S5**. Statistical analysis was done using both the one-sample *t*-test and paired *t*-test. Highest significance from either test is shown. *p*-values for both the tests are reported in **Table S2.** *\*p < 0.05, ** < 0.01*.

### miRNA regulate O-GlcNAcylation levels through OGT without impacting OGA

To assess the functional consequence of miRNA mediated OGT regulation, we investigated the impact of miRNA on the resultant O-GlcNAcylation levels. O-GlcNAc is controlled by both OGT and OGA levels. Thus, we tested the impact of a subset of miRNA mimics (down-miRs: let-7a-2-3p, miR-7-5p; up-miRs: miR-20a-3p, -148a-3p, -302b-3p) and NTC on OGT, OGA and O-GlcNAc in A549 cells using the same lysate for all three Westerns (**Fig. 5, Fig. S5, Table S2**). In this coupled dataset, we observed the expected down- and up-regulation of OGT although, in line with our previous data, we see some variability in the levels of response (**Fig. 5A-B, Fig. S5A-B**). In examining the impact of miRNA mimics on OGA, we noted that the NTC had an inhibitory effect on OGA expression when compared to untreated lysate (**Fig. 5C-D, Fig. S5C-D**). In contrast, NTC did not impact OGT levels (**Fig. S3E-F**). The NTC used is a microRNA from C. *elegans* which has been shown to impact the expression of other proteins (26, 28, 29). Hence, we used untreated cell lysate as the control for miRNA mimics when examining OGA levels. In general, we observed no changes in OGA expression with the tested miRNAs (**Fig. 5C-D, Fig. S5C-D**). However, we observed changes in O-GlcNAc levels that closely aligned with the alterations in OGT (fold changes OGT, OGA, O-GlcNAc, respectively: let-7a-2-3p: 0.45, 0.9, 0.54; miR-7-5p: 0.72, 0.76, 0.69; miR-20a-3p: 1.3, 0.96, 1.6; miR-148a-3p: 1.4, 1.0, 1.6; miR-302b-3p: 1.5, 0.93, 1.3, **Fig. 5, Fig. S5**). Thus, our data argues that these miRNAs regulate O-GlcNAcylation via direct alteration of OGT expression.

### OGT and O-GlcNAc levels are regulated by endogenous miRNA

To confirm that our miRNA mimics accurately represent the activity of endogenous miRNA, we utilized anti-miRs (miRNA hairpin inhibitors) (**Fig. 6, Fig. S6, Table S2**). These inhibitors bind to endogenous miRNA in AGO2 complexes, displacing mRNA and resulting in enhancement of protein levels for anti-down-miRs and repression of protein expression for anti-up-miRs. For the inhibitors to function, the targeted miRNAs must be expressed in the cell line. In A549, only miR-7-5p and -148a-3p were expressed at moderate levels, therefore we focused on these miRNAs. Although we failed to see statistically significant changes with either anti-miR due to variability in response, we observed a clear decrease of OGT expression with the anti up-miR-148a-3p (0.61 fold) and a slight increase with the anti-down-miR-7-5p (1.1 fold, **Fig. 6A-B, Fig. S6A-B**). We also analyzed the lysates to determine the impact of endogenous miRNA on O-GlcNAc levels. We observed a significant loss of O-GlcNAc expression upon treatment with anti-up-miR-148a-3p consistent with the loss observed for OGT (0.57 fold, Fig. **6C-D**, **Fig. S6C-D, Table S2**). In contrast, the anti-down-miR-7-5p did not impact O-GlcNAc levels.

**Figure 6.**
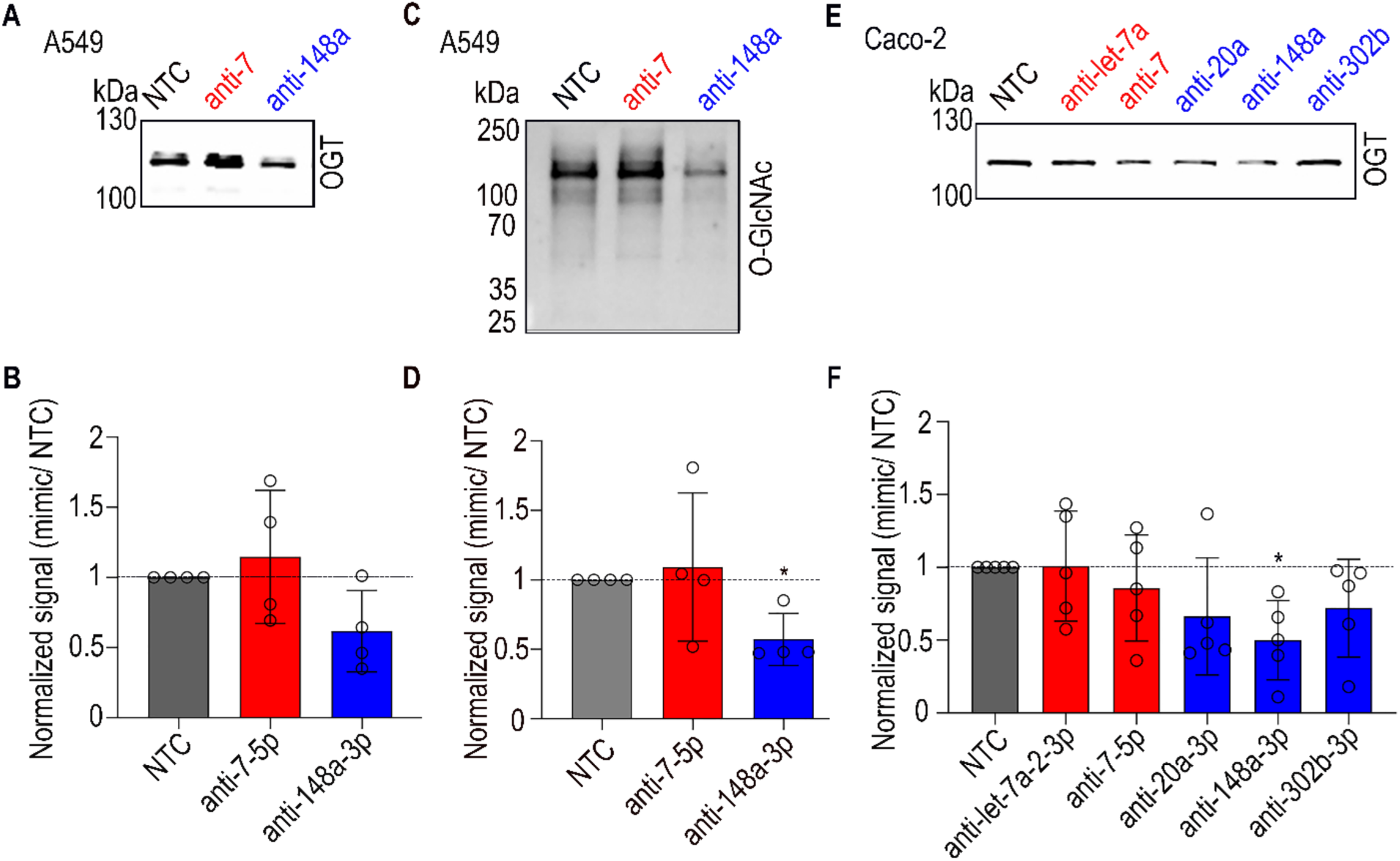
Inhibition of endogenous miRNAs impact OGT levels. (A, C, E) Representative Western blot analysis for α-OGT (A) and α-O-GlcNAc (C) in A549 and α-OGT (E) in Caco-2. Cells were transfected with 50 nM of either anti-miR non-targeting control (NTC, grey) or anti-miRs (anti-down-miRs (red): let-7a-3p, -miR-7-5p. anti-up-miRs (blue): -20a-3p, -148a-3p, -302b-3p) for 72 h prior to lysis and analysis. (B, D, F) Quantification of Western blot analysis for OGT (A, n=4), O-GlcNAC (C, n=4) in A549 or OGT (E, n=5) in Caco-2 cells. Bar charts represent average data for indicated biological replicates. For all quantification, Western blot samples were normalized first to total protein (Ponceau) and then to the normalized NTC. Individual replicates are indicated as discrete points. Error bars represent standard deviations. Additional associated data is shown in **Fig. S6**. Statistical analysis was done using both the one-sample *t*-test and paired *t*-test. Highest significance from either test is shown*. p*-values for both the tests are reported in **Table S2.** \**p < 0.05*

We next tested inhibition of endogenous miRNAs in CaCo-2. This cell line has very high expression of miR-148a-3p, moderate expression of miR-7-5p, low expression of miRs-302b-3p, and -20a-3p, and no detectable let-7a-2-3p (60). In line with the expression data, only inhibition of miR-148a-3p showed a strong significant response, with a clear loss of OGT expression in all replicates (0.49 fold, **Fig. 6E-F, Fig. S6E-F**). Some impact was also seen upon inhibition of miRs-20a-3p (0.66 fold) and -302b-3p (0.72 fold), in line with expression levels. The anti-miR for the known miRNA downregulator of OGT, miR-7-5p, did not show any impact in CaCo-2 despite expression of the miRNA. Overall, our results provide strong evidence that endogenous miR-148a-3p upregulates OGT protein expression and that this directly impacts levels of O-GlcNAc.

### Downregulation of OGT by the let-7 family is dependent on seed sequence conservation

The let-7 family was one of the earliest and largest families of miRNA to be discovered and is conserved from invertebrates to humans (61). In C. *elegans*, let-7 controls the timing of terminal differentiation and is important in development (62, 63). Like OGT, the let-7 family has a well documented role in the cell cycle, with high let-7 leading to cell cycle exit through targeting of multiple cell cycle genes.(64–66). In humans, there are 10 members of the let-7 family, with similar seed sequences (**Fig. 7**) (67, 68). With this is mind, we evaluated the impact of 8 members of the let-7 family on OGT expression by Western blot analysis (let-7a-2-3p, -7g-3p, -7c-3p, -7a-3p, -7b-3p, -7d-3p, -7e-3p, -7f-1-3p, -7f-2-3p, and -7i-3p, **Fig.7A-B, Fig. S7, Table S2**). We observed strong downregulation of OGT expression by let-7a-2-3p and -7g-3p (let-7a-2-3p: 0.3 fold, -7g-3p: 0.23 fold), consistent with the conserved seed site between the two (**Fig. 7C**). Significant downregulation of OGT was also observed in cells treated with let-7e-3p, -7f-2-3p and -7i-3p (let-7e-3p: 0.66 fold, -7f-2-3p 0.55 fold and -7i-3p 0.57 fold), but not other members of the family.

**Figure 7.**
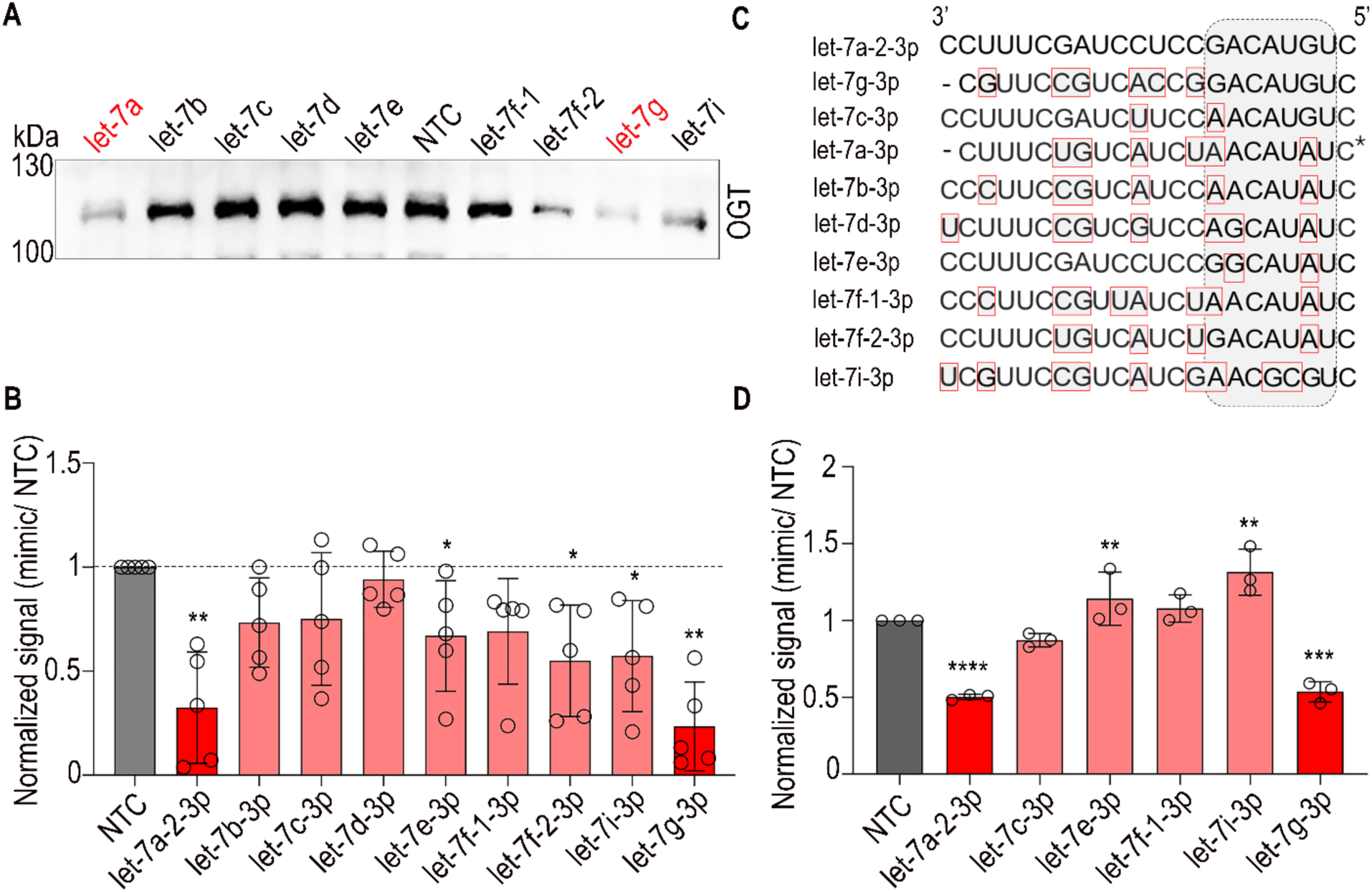
Let-7 family members downregulate OGT expression. (A) Representative Western blot analysis for α-OGT in A549 cells treated with 50nM of either NTC, (grey) or let-7 family members (let-7a-2-3p, -7b-3p, -7c-3p, -7d-3p, -7e-3p, -7f-1-3p, -7f-2-3p, -7g-3p and -7i-3p) for 72 h prior to lysis and analysis. (B) Quantification of Western blot analysis for OGT. Bar graphs represent average data from n=5 biological replicates. Data for let-7a-2-3p and -7g-3p are solid red for emphasis. Western blot samples were normalized first to total protein (Ponceau) and then to the normalized NTC. Individual replicates are indicated as discrete points. Error bars represent standard deviations. Additional associated data is shown in **Fig. S7**. Statistical analysis was done using both the one-sample *t*-test and paired *t*-test. Highest significance from either test is shown. *p*-values for both the tests are reported in **Table S2**. (C) Alignment of let-7 family members in order of seed similarity to let-7a-2-3p. The seed region is shown in grey box and differences between miRs are outlined in red. Let-7a-3p was not tested. (D) Small scale analysis of miRFluR assay for let-7 family. HEK-293T cells were co-transfected with pFmiR-OGT sensor and either NTC or select let-7 family members as indicated. Average signal (n=3 biological replicates) from miRFluR assay normalized over NTC is shown and individual replicates are indicated as discrete points. Error bars represent standard deviations. Statistics was done using paired *t*-test. *p*-values are reported in **Table S2.** \**p < 0.05, ** < 0.01, *** < 0.001*

Since let-7 family members share similar seed sequences, we aligned them in order of conservation of their seed region (**Fig. 7C**). As previously mentioned, let-7a-2-3p and -7g-3p have identical seeds and both show strong downregulation of OGT in both Western and using our miRFluR sensor (**Fig. 7D**, note: let-7g-3p was QCed out of the initial dataset). In contrast, changing a single base pair in the seed, as seen in let-7c-3p, abrogated the impact on OGT expression, and much of the signal in our miRFluR assay. Although let-7e-3p, -7f-1-3p and -7i-3p showed some downregulation of the OGT protein, no change was observed from 3’-UTR binding in either the original high-throughput or small scale miRFluR assay (**Figs. 1D & 7D**, **Dataset S1**), arguing that their impact is either indirect or through binding to the 5’UTR or coding region. Overall, our results are consistent with the importance of the seed region for let-7 family regulation of OGT via the 3’UTR and the impact that even a single nucleotide change can have.

To identify sites of let-7a-2-3p regulation in the 3’-UTR, we surveyed Targetscan, which relies on seed pairing (23). Two binding sites were identified, neither of which were observed in CLIP data (**Fig. 8A-D**). We made three mutant sensors to test the predicted sites (MUT-A for site +1757-1779 bp, MUT-B for site +2866-2886 bp and MUT A + B for both the sites together) mutating the sites to the sequence of let-7g-3p. We analyzed miRNA regulation in the wildtype and mutant pFmiR-OGT sensors with let-7a-2-3p or - 7g-3p mimics in HEK-293T cells, comparing them to NTC (**Fig. 8E, Table S2**). Mutating either site relieved some of the repression from let-7a-2-3p or -7g-3p. However, mutating both sites showed a more profound effect, almost completely abrogating the repressive effects of the miRNAs. The multiple sites might help to explain the strong repression observed from these miRNAs on OGT expression in cells (**Figs. 4 & 7**).

**Figure 8.**
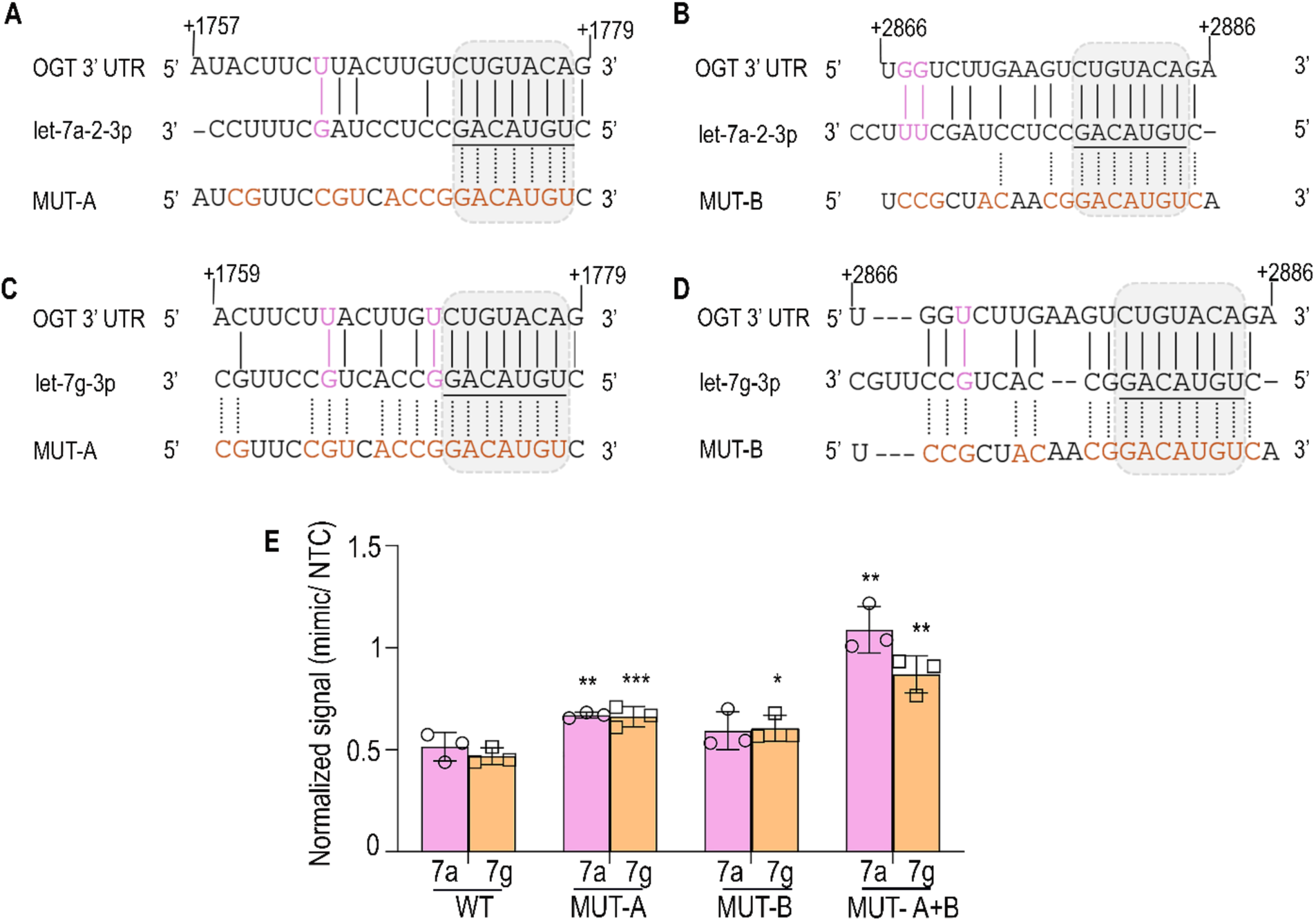
Validation of let-7a/g site on OGT 3’UTR. (A-D) Alignment of let-7a-2-3p and -7g-3p with their predicted OGT-3’ UTR sites and their corresponding mutants are shown. Mutated nucleotides (orange) and wobble base pairs (pink) are indicated. The miRNA seed sequence outlined in grey box and underlined. (E) Bar graph shows average data for HEK-293T cells transfected with let-7a-2-3p (pink) or -7g-3p (orange) with WT (wildtype) or MUT (Mutant-A, Mutant-B and Mutant A+B) miRFluR sensors. For each sensor, data was normalized over NTC in that sensor. The experiment was performed in biological triplicates. Statistics was done using paired *t*-test. *p*-values are reported in **Table S2**. For additional information see **Tables S3-5**. \**p < 0.05, ** < 0.01, *** < 0.001*.

### miR-148/152 family upregulates OGT expression via a direct AGO2 complex through multiple distributed binding sites

The upregulatory miRNA, miR-148a-3p belongs to the miR-148/152 family of miRNA which emerged in vertebrates and consists of miR-148a-3p, -148b-3p and -152-3p (**Fig. 9A**) (64, 69). These miRNA, which differ from one another by only 2-3 bases, are known to be involved in cancer and immune regulation (64, 70). All three of these miRNA were hits in our miRFluR assay (**Fig. 2D**) and were observed in CLIP datasets (53). Therefore, we tested whether they impacted OGT protein expression in A549 cells. In line with our previous observations for miR-148a-3p, both miR-148b-3p and -152-3p significantly upregulated OGT (∼2 fold, **Fig. 9B-C, Fig. S8A-B, Table S2**). Inhibition of endogenous miRNA with anti-miRs decreased OGT levels (anti-miR-148a-3p: 0.63 fold, -148b-3p: 0.88 fold, -152-3p: 0.86 fold, **Fig. 9D-E, Fig. S8C-D, Table S2**) with the strongest impact observed for anti-148a-3p, whose cognate miRNA has the highest expression in A549. We also observed a significant decrease in O-GlcNAc levels upon inhibition of all three endogenous miRNA (anti-miR-148a-3p: 0.50 fold, -148b-3p: 0.83 fold, -152-3p: 0.65 fold, **Fig. 9F-G, Fig. S8E-F, Table S2**), arguing for a direct impact of these miRNAs on O-GlcNAcylation via their regulation of OGT (**Fig. 9D-G, Fig. S8C-F, Table S2**). Overall, the data support a role for the miR-148/152 family in O-GlcNAc regulation.

**Figure 9.**
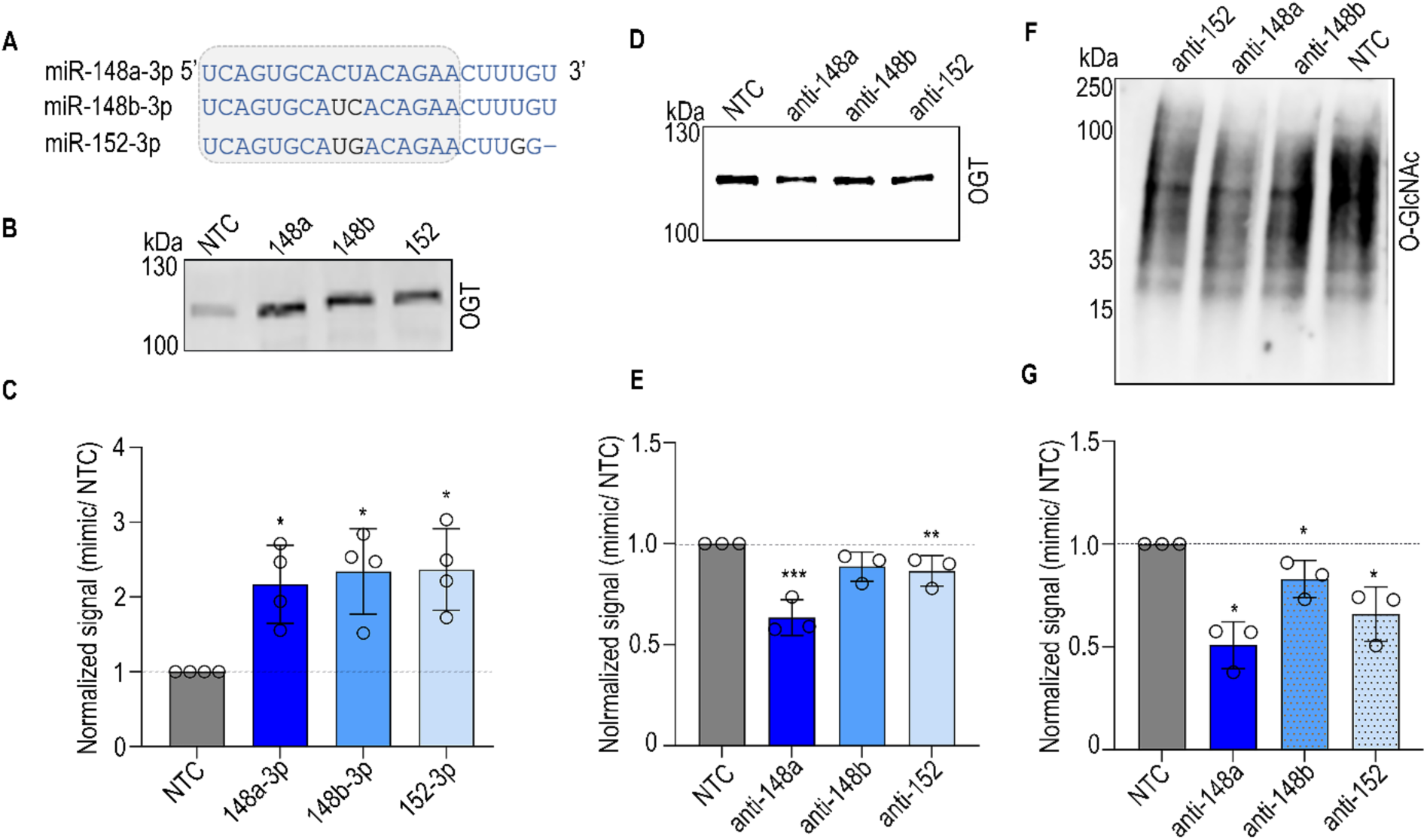
Endogenous miR-148/152 family upregulates OGT expression and O-GlcNAcylation. (A) Sequence alignment of miR-148/152 family. Conserved nucleotides are shown in blue. (B) Representative Western blot analysis for α-OGT in A549. Cells were treated with 50nM of either NTC (grey) or miR-148/152 family members (blue) for 72 h prior to lysis and analysis. (C) Quantification of Western blot analysis for OGT as in (B, n=4). (D, F) Representative Western blot analysis for α-OGT (D) and α-O-GlcNAc (F) in A549 cells treated with 50nM of either anti-miR non-targeting control (NTC, grey) or anti-up-miR-148/152 family members (blue) for 72 h prior to lysis and analysis. (E, G) Quantification of Western blot analysis for OGT (E) and O-GlcNAc (G) in A549 (n=3). Bar graphs represent average data from indicated biological replicates. Western blot samples were normalized first to total protein (Ponceau) and then to the normalized NTC. Individual replicates are indicated as discrete points. Error bars represent standard deviations. Additional associated data is shown in **Fig. S8**. Statistical analysis was done using both the one-sample *t*-test and paired *t*-test. Highest significance from either test is shown. Highest significance from either test is shown. *p*-values are reported in **Table S2**. *\*p < 0.05, ** < 0.01*.

CLIP datasets reporting potential interactions between OGT and the miR-148/152 family identified them in AGO2 complexes (53). In previous work, upregulation was shown to require AGO2 (29, 32, 33). AGO2 is also an important part of downregulatory complexes with miRNA (71). Thus, we tested whether regulation of OGT by either the down-miR let-7a-2-3p or the up-miR 148a-3p required AGO2. Silencing of AGO2 in A549 cells using pooled siRNA (NTP or siAGO2) for 48 h prior to transfection with miRNA caused a loss of both downregulation by let-7a-2-3p and upregulation by miR-148a-3p, reconfirming the importance of AGO2 complexes in miRNA-mediated regulation (**Fig. 10A-B, Fig. S9A-B, Table S2**). Based on the CLIP analysis, we anticipated that miR-148a-3p and the OGT mRNA are in a direct complex within AGO2. To confirm the presence of this complex, we overexpressed eGFP-AGO2 in an AGO-2 knockout HEK-293T cell line and transfected with either biotinylated NTC or miR-148a-3p mimic. Pulldown of the biotinylated miRNA showed AGO2 more strongly associated with miR-148a-3p, in line with expectations (**Fig. 10C, Fig. S9D-E**). RT-PCR analysis on equal amounts of RNA from the pulldowns showed a strong enrichment of OGT mRNA when miR-148a-3p mimic was used in comparison to the NTC, confirming the presence of a direct AGO2:*OGT*:miR-148a-3p complex (**Fig. 10D, Table S2**).

**Figure 10:**
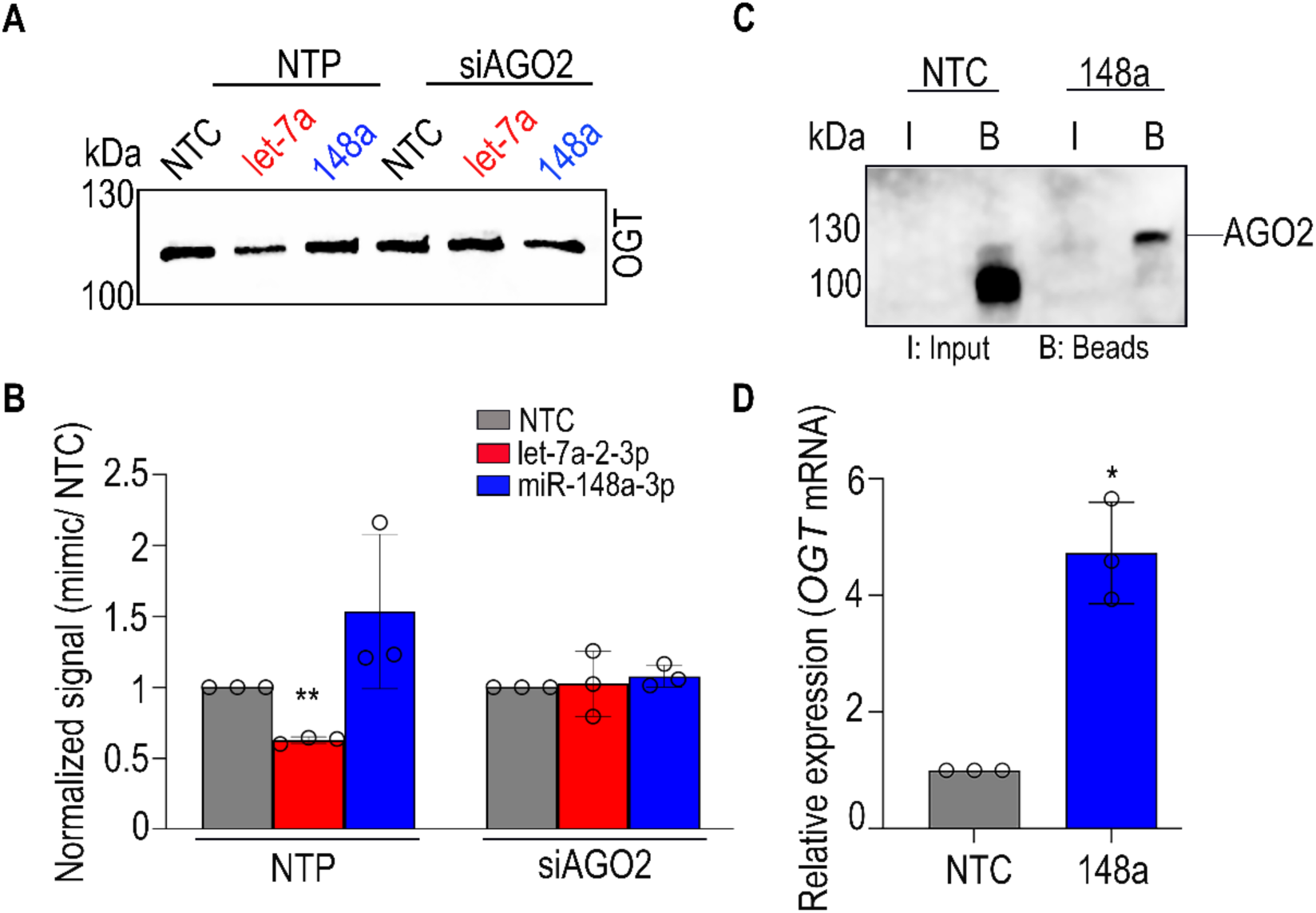
Upregulation of OGT by miRNAs requires direct interaction with its 3′UTR within a complex containing AGO2. (A) Representative Western blot of α-OGT in A549 cells treated with 50nM of either NTC, let-7a-2-3p (down-miR) or miR-148a (up-miR) for 72 h post transfection of these cells with 20nM of pooled siRNA (non-targeting pool, NTP or si-AGO2) for 48 h. (B) Quantification of Western blot analysis as in (A). n=3 biological replicates. For all quantification, Western blot samples were normalized first to total protein (Ponceau) and then to the normalized siNTC/NTP. Individual replicates are indicated as discrete points. Error bars represent standard deviations. Additional associated data is shown in **Fig. S9**. Statistical analysis was done using both the one-sample *t*-test and paired *t*-test. Highest significance from either test is shown. (C) Pulldown of α-AGO2 by biotinylated miR-148a-3p. eGFP-AGO2 cells were transfected with biotin labelled NTC or miR-148a-3p (50nM) for 72 h prior to lysis and immunoprecipitation. Complexes were resolved by SDS-PAGE and analyzed by Western blot analysis using an α-AGO2 antibody (n=3 biological replicates). Input represents 1% of total lysate used for immunoprecipitation. Representative Western blot is shown. (D) Quantification of RT-qPCR of OGT. RNA was isolated from pulldown material as in (C). Additional information is in **Table S3**. Graph shows normalized OGT mRNA expression relative to GAPDH, average of biological replicates is shown and individual points are indicated (n=3). All *p*-values are reported in **Table S2.** \**p < 0.05*.

We next sought to determine binding sites for the miR-148/152 family on the 3’UTR of OGT. CLIP analysis identified two AGO2 binding sites for these miRNA. The annotation of CLIP data generally relies on seed-dependent algorithms, thus many upregulatory sites are not properly annotated in these datasets as they are not seen by the computational models currently used (53, 54). Despite this, one canonical and one non-canonical CLIP site was identified for the interaction of miR-148a-3p with OGT (**Tables S6 & S7,** sites 1-2). RNAhybrid analysis identified the non-canonical site as the strongest predicted binding site for the miR-148/152 family (72). This site was also identified by the miRanda algorithm (73). Mutation of this site, however, gave only a small, non-statistical loss of upregulation for all family members (site 1, Normalized miR-148a-3p/NTC; WT: 1.6 fold, single mut. 1.46 fold, **Fig. 11A-C, Fig. S10A,C**). We next generated a double mutant that ablated both CLIP sites and found no additional change in upregulation (sites 1,2; double mut: 1.49 fold, **Fig. 11A-C. Fig. S10A,C**). Analysis by RNAhybrid of potential binding sites for miR-148a-3p identified 5 more sites distributed throughout the 3’UTR with minimum free energy values (mfes) <-19 kcal/mol (sites 3-7, **Fig. 11A-B**, **Table S7**), within the range of previously identified upregulatory sites (27, 74). To test the importance of the additional sites, we created a series of truncation mutants (**Fig. 11B**, T1: ∼1500bp, T2: ∼1Kb and T3: ∼500bp). We used the empty pFmiR vector, which does not contain a 3’UTR, as a negative control. For all three truncations, we observed partial loss of upregulation relative to the full-length construct (Normalized miR-148a-3p/NTC; WT: 1.60 fold, T1: 1.35 fold, T2: 1.32 fold, T3: 1.25 fold, **Fig. 11D**), although only T2 and T3 were statistically significant. Of note, although T3 was not statistically different from the empty vector (Empty: 1.07 fold), it did show some upregulation, arguing that sites 6 and 7 might still contribute., Similar data is seen for the other two members of the family, miR-148b and miR-152 (**Fig. S10B, D**, **Tables S8 & S9**). Our data supports a model in which OGT upregulation by the miR-148/152 family is via multiple binding sites distributed through the OGT 3’UTR.

**Figure 11:**
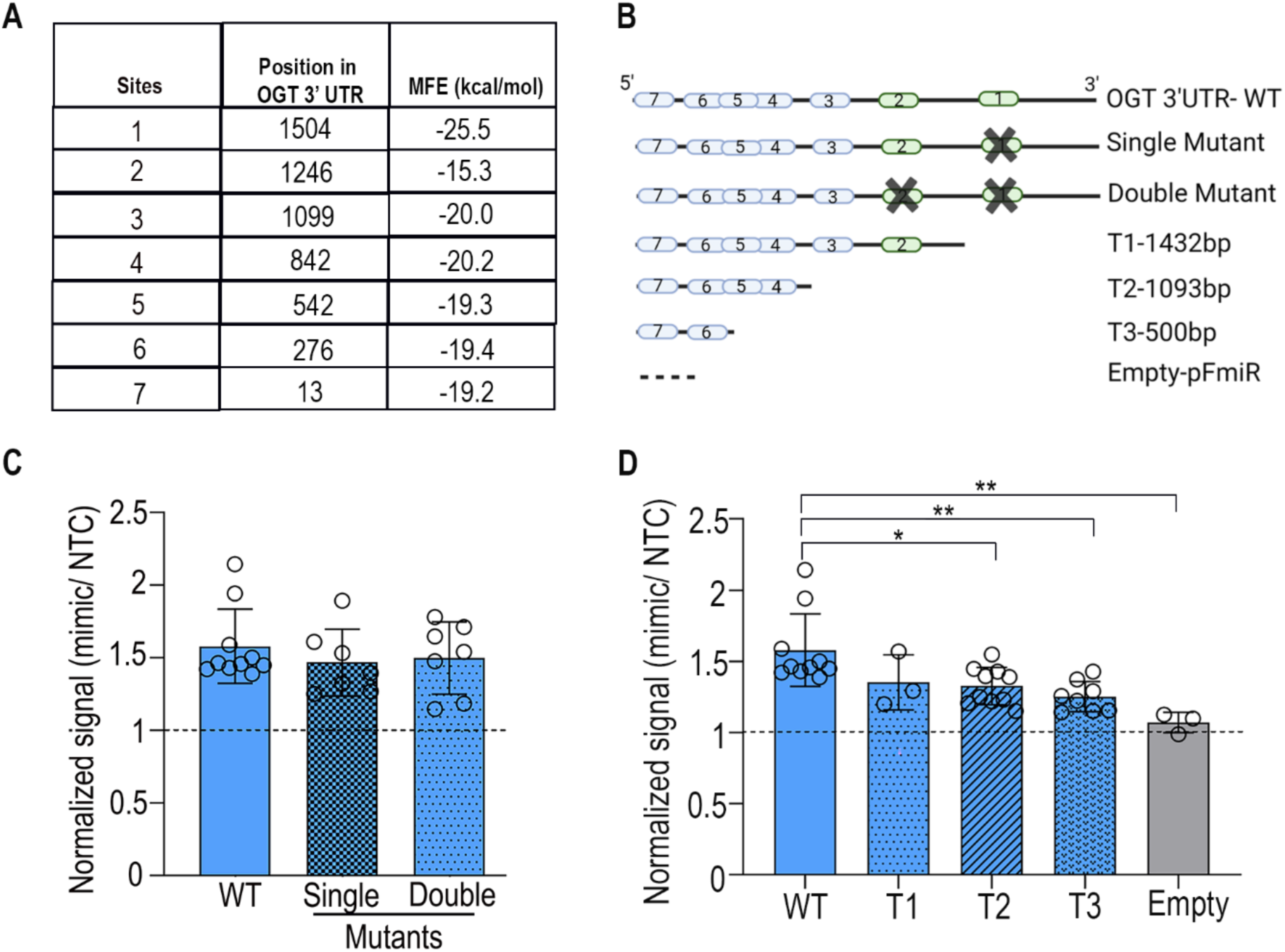
miR-148a upregulates OGT through multiple binding sites. (A) miR-148a binding sites identified in CLIP analysis (sites 1-2) or predicted by RNAhybrid (sites 1, 3-7), with their position in the OGT 3’UTR and respective minimum free energy (mfe) indicated. (B) Schematic representation of OGT 3’ UTR mutants and truncations.CLIP sites are represented in green; additional RNAhybrid binding sites are shown in blue. (C) Bar graph shows average data for HEK-293T cells transfected with miR-148a-3p (with WT (wildtype) or MUT (Single, Double) miRFluR sensors. (D) Bar graph shows average data for HEK-293T cells transfected with miR-148a-3p (with WT (wildtype), truncated (T1, T2, T3) or empty pFmiR) miRFluR sensors. For all graphs, data was normalized over NTC in that sensor (n≥3 biological replicates, individual replicates indicated). Statistics were done using two-way ANOVA with Tukey’s comparison. *p*-values are reported in **Table S2**. \**p < 0.05, ** < 0.01*.

## Discussion

O-GlcNAc transferase (OGT) is a dynamic regulatory hub that responds to a variety of cellular signals including metabolism and stress (46, 75, 76). Control over OGT levels, and the resultant levels of O-GlcNAc, requires a high degree of transcriptional and post-transcriptional regulation of the enzyme, including by miRNA (37–42, 49, 50, 77–80). Although traditionally thought to downregulate, it has become clear that the impact of miRNA is bidirectional and that upregulation of protein expression is a common occurrence (26–29, 31, 33, 34). Herein we identify ∼200 miRNA regulators of OGT, including 59 down-miRs and 125 up-miRs, using our high-throughput miRFluR assay (26–30) (**Fig. 2, Table S1, Dataset S1**). Our hit list included 3 known down-miRs and AGO2 binding sites for 32 of our hits (17 down-miRs, 15 up-miRs) were previously identified in CLIP data (**Datasets S1 & S2**). In line with previous work, data from our miRFluR assay accurately identified the impact of miRNA on protein levels, in this case on OGT (26–30). Although our validation experiments were generally consistent, we did find that OGT levels showed higher variability in response to miRNA than other genes, perhaps due to the the high sensitivity of OGT to various forms of stimuli including glucose levels and stress (1, 46) (81). We also observed strong cell dependent effects for some miRNA, including the down-miR miR-7-5p, a known OGT regulator validated in multiple studies (40, 55, 82). This is in line with the cell line dependency observed for other miRNA regulatory interactions (**Figs. 3 & 4**) (26, 28, 29, 57). In mapping the OGT regulatory landscape, our work affirms the bidirectionality of miRNA regulation of OGT.

The regulation of O-GlcNAc is complex, responding to OGT and OGA levels as well as changes in the hexosamine biosynthetic pathway, thus it was unclear whether miRNA that regulated OGT would have a direct impact on O-GlcNAc levels (13),(46),(11). There is also strong evidence that the levels of OGA and OGT can be coupled (81, 83, 84). In this work, we found that the miRNA impacting OGT did not impact OGA and yet directly affected O-GlcNAc levels in line with their effects on OGT (**Figs. 5 & 9**). It should be noted however that we only tested a subset of the miRNA from our hitlist, thus it is possible that other miRNA may co-regulate the two genes. Analysis of this will require mapping of the miRNA regulation of OGA, a subject of future work.

OGT is a highly conserved gene (17–19). One third of miRNA families are highly conserved across species (85), with 60% of miRNA loci conserved between humans and mice (86). The let-7 family, several members of which inhibited OGT in our assay, is one of the earliest miRNA families to emerge in evolution. Both let-7a-2-3p and let-7g-3p inhibited OGT expression through seed-dependent binding (**Figs. 7 & 8**). Strong regulation by these miRNAs required two canonical binding sites, one at chromosomal position chrX: 70,793,730 (1771 site) and the other at position chrX: 70,794,824 (2878 site) within the human 3’UTR for OGT. Analysis of their conservation found that the 1771 site emerged in placental mammals, with evidence for it in elephants, but not in opossum (**Table S4**). The 2878 site, however, is far more recent, with evidence for it in primates, but not other mammals (**Table S5**). Tight regulation of OGT expression required both binding sites (**Figs. 7 & 8**). This binding interaction may be related to the roles of both OGT and let-7 in human health, most notably type 2 diabetes which emerged in primates and thus is an evolutionarily recent disease (87).

Both, OGT and let-7a are known to play key roles in the metabolic pathways underlying diabetes. However, the precise role of OGT and O-GlcNAc in formation of type-II diabetes is still unclear (88, 89). Deletion of OGT in islet β-cells causes severe hyperglycemia, glucose intolerance and severe diabetes in mice (14) and O-GlcNAc positively regulates the secretion of insulin (90, 91). O-GlcNAc levels are closely tied to glucose levels and may also play a role in downstream damage from diabetes (88, 89) (92, 93). The let-7 family, including let-7a/g is known to regulate multiple proteins underlying glucose metabolism, including the insulin receptor (94). Knockdown of let-7 improved insulin resistance in obese mice (95). Although the importance of let-7a/g regulation of OGT in glucose metabolism is not known, the evolutionary appearance of the site and the regulatory roles of both the miRNA and the gene are intriguing.

Unlike the let-7 family, the miR-148/152 family emerged later in evolution, first appearing in jawless fish (96). All three members of the miR-148/152 family (miR-148a-3p, -148b-3p and -152-3p) upregulated OGT expression. In line with this, analysis of TCGA datasets using miRactDB identified positive correlations between miR-148a-3p and OGT in two cancers (head & neck and prostate) and in three adjacent tissues (breast, kidney chromophobe, kidney renal clear cell carcinoma) (97). Regulation by miR-148a-3p required AGO2, which complexed with the miRNA and OGT mRNA in cells (**Fig. 10**). Unlike let-7a/g-3p, two miR-148/152 binding sites were identified in CLIP pulldowns of AGO2 (**Table S6, Dataset S2**). However, ablation of these sites had only a small impact on upregulation by the miR-148/152 family, implying that multiple sites might exist (**Figs. 11 & S10**). Most miRNA are thought to regulate via canonical seed interactions and the algorithms predicting miRNA interactions are biased towards that assumption (23). To date, the majority of experimentally validated upregulatory miRNA binding sites identified through our high-throughput approach are non-canonical, providing an explanation for why upregulation by miRNA has generally been overlooked (26–29, 74). RNAhybrid analysis of the OGT 3’UTR revealed 5 more potential non-canonical sites with mfes within range of previously identified upregulatory sites (**Tables S7-9**). Analysis of truncated 3’UTRs, in which sites 1, 1-3, or 1-5 were deleted, showed a small but clear decrease as more sites were removed, indicating that miR-148/152 requires multiple sites for upregulation that are distributed across the OGT 3’UTR. A similar phenomenon has been observed for downregulation of HMGA2 by let-7, where 6 independent sites were identified across the 3 kb 3’UTR (98).

Previous work hints at multiple mechanisms for upregulation which may involve distinct protein complexes, including impacts on both mRNA stability and translation (26, 28, 29, 99) (33). Although the precise mechanism of upregulation by miRNA is still a mystery, our data confirms the requirement for AGO2, the existence of a direct non-canonical miRNA:mRNA complex and the presence of multiple upregulatory binding sites for upregulation of OGT by miR-148a-3p. Taken together, our data provides important insights into OGT and its regulation by miRNA.

## Experimental Procedures

### Cloning

OGT 3’ UTR (ENST00000373719.3) was amplified from cDNA (HEK-293T cell line) using Q5 High-Fidelity DNA Polymerase (Catalog # M0492L, New England Biolabs (NEB)) according to the PCR conditions provided by NEB, using the primers shown in **Table S3**. NheI and BamHI restriction sites were used for cloning the OGT 3’ UTR downstream from Cerulean in the pFmiR-empty backbone (30) using the standard NEB ligation protocol to create pFmiR-OGT. The cloned sequence was verified by Sanger Sequencing (Molecular Biology Services Unit, University of Alberta). NucleoBond Xtra Maxi EF (Ref. 740424.50, Macherey-Nagel) was used for large scale endotoxin free DNA preparation of the verified plasmid. The plasmid map for pFmiR-OGT and its corresponding 3’ UTR sequence can be found in **Figs. S1 & S2**.

### Cell lines

All cell lines (HEK-293T, A549, CaCo-2 and SH-SY5Y) with the exception of the HEK-293T-AGO2 knockout line were purchased from the American Type Culture Collection (ATCC) and cultured in suggested media; HEK-293T in Dulbecco’s Modified Eagle medium (DMEM) supplemented with 10% Fetal Bovine Serum (FBS); A549 in FK-12 medium supplemented with 10% FBS, CaCo-2 in DMEM supplemented with 20% FBS and SH-SY5Y in DMEM supplemented with 1% Non essential amino acids (NEAA) and 10% FBS. The HEK-293T-Ago2KO cell line was purchased from GenScript and cultured in the same media as HEK-293T. The biological sex of the cell lines used is as follows: female: HEK-293T and SH-SY5Y, male: CaCo-2 and A549 (100). All the cells were maintained under standard conditions (5% CO2 at 37°C) and used below passage 20.

### Stable cell line development (HEK-293T-eGFPAGO2)

The plasmid, pHAGE-eGFP-Ago2wt, was purchased from Addgene (Cat. 74231). Lentivirus particles were prepared using third generation packaging mix (Applied Biological Materials, Cat. LV053) following the manufacturers protocol using the HEK-293T Lenti-X cell line. A series of dilutions ranging from 1:2 – 1:100 was used for transduction of HEK-293T-Ago2KO cells following the online Addgene protocol (https://www.addgene.org/protocols/generating-stable-cell-lines/). Cultures with the highest percentage of GFP expression by microscopy were harvested between days 14-18 post-transduction and sorted for GFP positive cells. The presence of eGFP-AGO2 in the cells was verified by Western blot analysis.

### miRFluR high-throughput assay

The Human miRIDIAN miRNA mimic library version 21.0 (Dharmacon, v. 21) was resuspended in ultrapure nuclease-free water (REF. #: 10977-015, Invitrogen) and aliquoted into black 384-well flat clear bottom black tissue culture-treated microplates (Product Number: 3764, Corning). Each miRNA was aliquoted in triplicate (2 pmol/well) together with six wells of Non-Targeting Control (NTC, CN-001000-01, Horizon Discovery) in each plate. To each well was added 25 ng of pFmiR-OGT sensors diluted in 5µL of Opti-MEM (Gibco) and 0.1 µL of Lipofectamine 2000 Transfection Reagent (Catalog# 11668500, Life Technologies) diluted in 5 µL of Opti-MEM (pre-incubated at room temperature for 5 min). The plates were then spun at 300 rpm for 30 s using Centrifuge 5430 (Eppendorf) and incubated at room temperature for 20 min, after which 25 µL of HEK-293T (4 x 10^5^ cells/mL prepared in phenol red free DMEM supplemented with 10% FBS, 10-25µL/10mL penicillin-streptomycin) was added per well. After 48 h incubation at standard conditions (5% CO2 at 37 °C) the fluorescence signals for Cerulean (excitation (ex): 433 nm; emission (em): 475 nm) and mCherry (ex: 587 nm; em: 610nm) were measured using a Synergy H1 microplate reader (BioTek) with the Gen5 software (version 3.08.01) at 48 h. The entire assay was done in duplicate in two independent experiments.

### Data analysis of high-throughput miRFluR assay

The ratio of the fluorescence signal for Cerulean and mCherry (Cer/mCh) was calculated for each well in an individual plate. For each miRNA, the values of the ratios were averaged across wells and the standard deviation (S.D.) calculated. The % error of measurement (100 x S.D./mean) was then calculated for each miRNA. Plates where the median % error was >15% were removed. We also removed miRNA with >15% error. As we ran the assay in duplicate, we averaged the values for miRNA that passed QC in both datasets. We included miRNA that passed QC in only one dataset, using only data from the dataset where it passed QC. Post QC, this resulted in 2214 miRNAs of the 2601 miRNAs screened in our final list (85% of library). The mean Cer/mCh ratio for individual miRNA was normalised to the mean Cer/mCh ratio for the NTC within the same plate and S.D. were used to propagate the error. The data was then z-scored and the z-score (±1.65), corresponding to a *two*-tailed *p*-value of 0.10 (i.e. 90% confidence interval), was used as a threshold for significance. Post-analysis we identified 184 miRNA hits for OGT within 90% confidence interval **(see Fig. 2B-D, Table S2 and Dataset S1).**

### Generation of lysates for Western blot analysis

Western blot analysis was conducted on lysates from three cell lines: A549, CaCo-2 and SH-SY5Y. Cells were seeded into six-well tissue culture treated plates (A549 & CaCo-2: 3 x 10^5^ cells per well, SH-SY5Y 5 x 10^5^ cells) and cultured for at least 24 h in their respective media. Media was then removed and miRNA mimic or anti-miR was added. For miRNA mimic transfection, cells were transfected with miRNA mimics or non-targeting control (50 nM mimic Dharmacon, Horizon Discovery, 5 µl Lipofectamine 2000, Life Technologies in 250µl OptiMEM (Gibco)). For anti-miR transfections, cells were transfected with either miRIDIAN microRNA hairpin inhibitor negative control or miRIDIAN microRNA hairpin inhibitors anti-miRs: (-let-7a-2-3p, -miR-7-5p, -20a-3p, - 148a-3p, -302b-3p, 20 nM) as described above for miRNA mimics. After 16 h, media was changed to standard media. Cells were then harvested 72 h after transfection. Cells were washed 1 x with ice cold Hanks buffered salt solution (HBSS, Gibco) on ice and then lysed in RIPA lysis buffer (∼300-600 μL, Thermo Scientific) supplemented with protease inhibitor (cat. # 78446, Thermofisher). Cell lysate was transferred to an eppendorf tube and incubated on ice for 20 min, followed by centrifugation at 14,000 rpm for 15 min at 4 °C. The supernatant was transferred to a fresh tube and the protein concentration quantified by the Bradford Protein Assay (BioRad). Lysates were used for Western blot analysis as described below.

### Testing the impact of AGO2 knockdown on miRNA regulation

A549 cells (2 x 10^5^) were seeded into six well plates (Corning) and cultured for ∼24 h under standard conditions. Cells were then transfected with either 20 nM of pooled siAGO2 (SMARTpool, Dharmacon) or control (NTP, Dharmacon) using Lipofectamine™ RNAiMAX transfection reagent (catalog #: 13778150, Thermofisher) in OptiMEM following the manufacturer’s instructions. After 48 h, cells were transfected with miRNA mimics (NTC, -let-7a-2-3p and miR-148a-3p) as described previously. To help keep cells less stressed, the media was changed to standard media 16 h post-miRNA transfection. Cells were then lysed and analysed 72 h post-transfection of miRNA mimics as described above. Inhibition of AGO2 by the pooled siAGO2 (Dharmacon) was validated by RT-qPCR for this experiment and was previously validated by Western blot (29).

### Western blot of OGT, OGA, O-GlcNAc and AGO2

For all Western blot analysis, we ran 20 μg of protein. Protein was mixed with Laemeli buffer with 5 mM beta-mercaptoethanol and denatured for 10 min 100 °C prior to being run on SDS-PAGE gels using standard conditions. For OGT/OGA 10% SDS-PAGE gels were used, for O-GlcNAc 8% gels, and for AGO2 4-12% gradient gels. Proteins were then transferred to nitrocellulose (Invitrogen) using the iBlot2 transfer device (Invitrogen) and associated protocols. Membranes were then incubated with Ponceau S solution (Boston BioProducts, catalog #ST-180) for 5 min and the whole blot was imaged using protein gel mode (Azure 600, Azure Biosystems Inc.) to determine total protein levels per lane. Ponceau staining was removed by washing the membrane with Tris buffered saline (TBS, pH 8). Blots were blocked using 5% non-fat dry milk in TBST buffer (TBS with 0.1% Tween 20) for 1 h at room temperature with rocking (60 rpm, LSE platform rocker, Corning). The blocking buffer was removed and blots were rinsed briefly with TBS. Blots were then incubated with either α-OGT antibody (Abcam, cat # ab96718, 1:1000 in 5% BSA in TBST), α-O-GlcNAc antibody (RL2, Abcam, cat # ab2739 1:1000 in 5% BSA in TBST) or α-OGA (Thermo, cat # 61425, 1:1000 in 5% BSA in TBST) or α-AGO2 (Sigma, cat #ZRB1205, 1:1000 in 5% skim milk in TBST) overnight at 4°C with rocking. We validated the α-OGT and α-OGA antibodies using pooled siOGT and siOGA, respectively in A549 cells (Horizon Discovery, **Figs. S3C-D & S6G-H**). After an overnight incubation, the blots were washed 5 x for 2 min each with 0.1% TBST buffer on a reciprocating shaker. Membranes were then incubated with 1:10000 dilution of the appropriate secondary antibody (O-GlcNAc: goat α-mouse IgG-HRP, Jackson laboratory; α-OGT, α-OGA, α-AGO2: α-rabbit IgG-HRP, Jackson laboratory) in 5% BSA in TBST for 1 h with shaking (60 rpm) at room temperature. Membranes were then washed 3 x with 0.1% TBST buffer and developed using Clarity Max Western ECL substrate according to the manufacturer’s protocol (Bio-Rad). Membranes were imaged using chemiluminescent mode (Azure 600, Azure Biosystems Inc.).

### Data analysis of Western blot

Western blots were quantified using ImageJ software (ImageJ 1.54g, Java 1.8.0_345) (101). For each lane, signal was normalized to the Ponceau for that lane. For each blot, the Ponceau normalized signal for miRNA mimics was divided by the Ponceau normalized signal for NTC to give the data shown in graphs corresponding to Western blot analysis. We tested for statistical significance using two different statistical tests: the one-sample *t*-test against NTC=1 and the paired *t*-test (**Table S2**). For the paired t-test, the samples were paired by experiment.

### Multi-Site Mutagenesis

The 3’-UTR sequence of OGT and the miRNA sequences (let-7a-2-3p, let-7g, miR-148a-3p, -148b and 152) were analyzed with the RNAhybrid tool, which calculates the minimal free energy of hybridization for the target RNA sequence and miRNA. The two most stable predicted miRNA: mRNA interaction sites were selected for downregulators and mutant pFmiR-sensors were designed. While for upregulators, the +two CLIP binding sites were selected and mutants were generated for the strongest CLIP binding sites (single mutation) or both CLIP sites (double mutation). Mutation sites were designed using the NEBChanger tool (version v2.5.1, New England Biolabs (NEB)) (**Table S3**), and primers were ordered from Integrated DNA Technologies (IDT). The pmiR-OGT 3’ sensor was mutated using Q5 site-Directed Mutagenesis Kit (cat # E0554S, NEB) using manufacturer’s protocols. Mutations were confirmed by whole plasmid sequencing (Plasmidsaurus). The mutant plasmids were then used in the miRFluR assay as described above for the high-throughput version. A minimum of 4 wells were transfected per sensor and the analysis was done in 3 independent experiments. Data was analyzed using 2-way ANOVA with Tukey’s comparison.

### Truncations of OGT 3’UTR

A series of 3’ deletion constructs were made for the 3’UTR of OGT, with the 3 ends terminating at bases +1432 (Truncation-T1), +1093 (T2) and +500 (T3). The numbering of bases is designated relative to nucleotide 1 in the full length 3’ UTR (**Fig. S2**). Multiple primers were designed (**Table S3**) and, ordered from Integrated DNA Technologies (IDT). NheI and BamHI restriction sites were used for cloning the truncated OGT 3’ UTR downstream from Cerulean in the pFmiR-empty backbone (30) using the standard NEB ligation protocol to create pFmiR-OGT T series. Truncation sequence and orientation was confirmed by whole plasmid sequencing (Plasmidsaurus). These truncated plasmids were then used in the miRFluR assay as described above for the high-throughput version. As a negative control, pFmiR empty (without OGT 3’UTR) was used. A minimum of 6 wells were transfected per sensor and the analysis was done in at least 3 independent experiments. Two-way ANOVA with Tukey’s comparison was used to compare sensor data.

### miRNA Pulldown of OGT mRNA

Cells (3 x 10^6^ HEK-293T-eGFPAGO2) cells were seeded in 10 cm plates and cultured for ∼24 h in their respective media. Cells were then transfected with either biotin-labelled NTC control or miR-148a-3p (50 nM, Dharmacon Horizon Discovery, Cambridge, UK) using Lipofectamine™ 2000 transfection reagent in OptiMEM following the manufacturer’s instructions (Life Technologies). After 16 h, media was changed to standard culture media and cells were lysed 72 h post-transfection using 1% NP-40 lysis buffer (1M Tris-HCl, 1M NaCl, 1% NP-40, 0.5M EDTA). For Ago2 pull down, 50 μL of streptavidin magnetic beads (Pierce, ThermoFischer, Catalogue #88817) were separated on a magnetic stand and the clear supernatant was removed. Beads were then washed 3 x times with 1% NP-40 lysis buffer, blocked with 200 μL of bead blocking buffer (1 µg/µL BSA and 1 µg/µL Yeast tRNA (Thermo Catalogue # AM7119) in lysis buffer) and incubated with rotation at 4 °C for 3 h. After blocking, the supernatant was separated on the magnet stand and beads were washed 3 x times with 1% NP-40 lysis buffer. Beads were then resuspended and incubated with 2 mg of protein at 4°C for 1 h with rotation. The supernatant was separated on the magnet stand and beads were washed 3 x times with 1% NP-40 lysis buffer. Beads were used for further analysis for both protein immunoprecipitation-Western blotting for AGO2 and isolation RNA using TRIzol^TM^ for *OGT* mRNA analysis using qRT-PCR.

### cDNA Synthesis and RT-qPCR

Total RNA was isolated from cells treated as in miRNA mimic (A549) or miRNA pulldown (HEK-293T-eGFPAGO2) experiments using TRIzol^TM^ reagent (cat #: 15596018, Invitrogen) according to the manufacturer’s instructions. RNA concentrations were measured using NanoDrop, and 1 µg of isolated total RNA was reverse-transcribed to cDNA using Superscript III first-strand cDNA synthesis kit (Invitrogen, cat #: 18080300) according to manufacturers protocol using OligodT primers. Reverse transcription quantitative PCR (RT-qPCR) was performed using the SYBR Green method and cycle threshold values (Ct) were obtained using an Applied Biosystem (ABI) 7500 Real-Time PCR machine. GAPDH served as the housekeeping gene for normalization. Three biological replicates were conducted for each experiment. The primer sequences used in RT-qPCR can be found in **Table S3**.

## Data availability

The authors declare that all data can be found in this document and its supporting files.

## Supporting Information

This article contains supporting information:

Figs. S1-S11

Tables S1-S9

Datasets S1-S2 (separate Excel files)

## Author contributions

Conceptualization: Z.A.S., C.T., L.K.M.; Methodology: Z.A.S., F.J.C., C.T., R.D., L.K.M.; Investigation: Z.A.S., H.N; Resources: Z.J.; Validation: Z.A.S., H.N.; Formal Analysis: Z.A.S., H.N., R. D., H.H., L.K.M.; Visualization: Z.A.S., H.N., L.K.M.; Writing-Original Draft: Z.A.S., L.K.M; Writing-review & editing: Z.A.S., D.M., L.K.M.; Supervision: D.M., L.K.M.

## Funding and additional information

Funding for L.K.M. comes from the Canada Excellence Research Chairs Program (CERC in Glycomics). FACS experiments were performed at the University of Alberta Faculty of Medicine & Dentistry Flow Cytometry Facility, RRID:SCR_019195, which receives financial support from the Faculty of Medicine & Dentistry and Canada Foundation for Innovation (CFI) awards to contributing investigators. Figs. 1 and 2 were made using BioRender.

## Conflict of Interest

The authors declare no competing interests.

## Abbreviations

The abbreviations used are as follows:

OGT: O-GlcNAc transferase;
OGA: O-GlcNAcase;
AGO2: Argonaute 2;
3’UTR: 3’-untranslated regions;
mRNA: messenger ribonucleic acid;
microRNA: miRNA,
miR: micro ribonucleic acid;
siRNA: small interfering RNA;
NTP: non-targeting control pool;
NTC: non-targeting control,
down-miR: downregulatory miRNA;
up-miR: upregulatory miRNA;
WT: wildtype;
MUT: mutant;
SDS-PAGE: sodium dodecyl sulfate polyacrylamide gel electrophoresis;
PCR: polymerase chain reaction,
RNA-IP: RNA Immunoprecipitation.

## Notes

### Competing Interest Statement

The authors have declared no competing interest.

### Summary of Updates

Due to an issue of sequencing, the site identified in the former Fig. 10 was not correct. We have added new data on the mIR-148 site, added data and Figures accordingly.

