## Supplemental for "Analyzing the miRNA regulatory landscape of OGT identifies bidirectional regulation"

**This file contains:**

**Figs. S1-S11**

**Tables S1-S9**

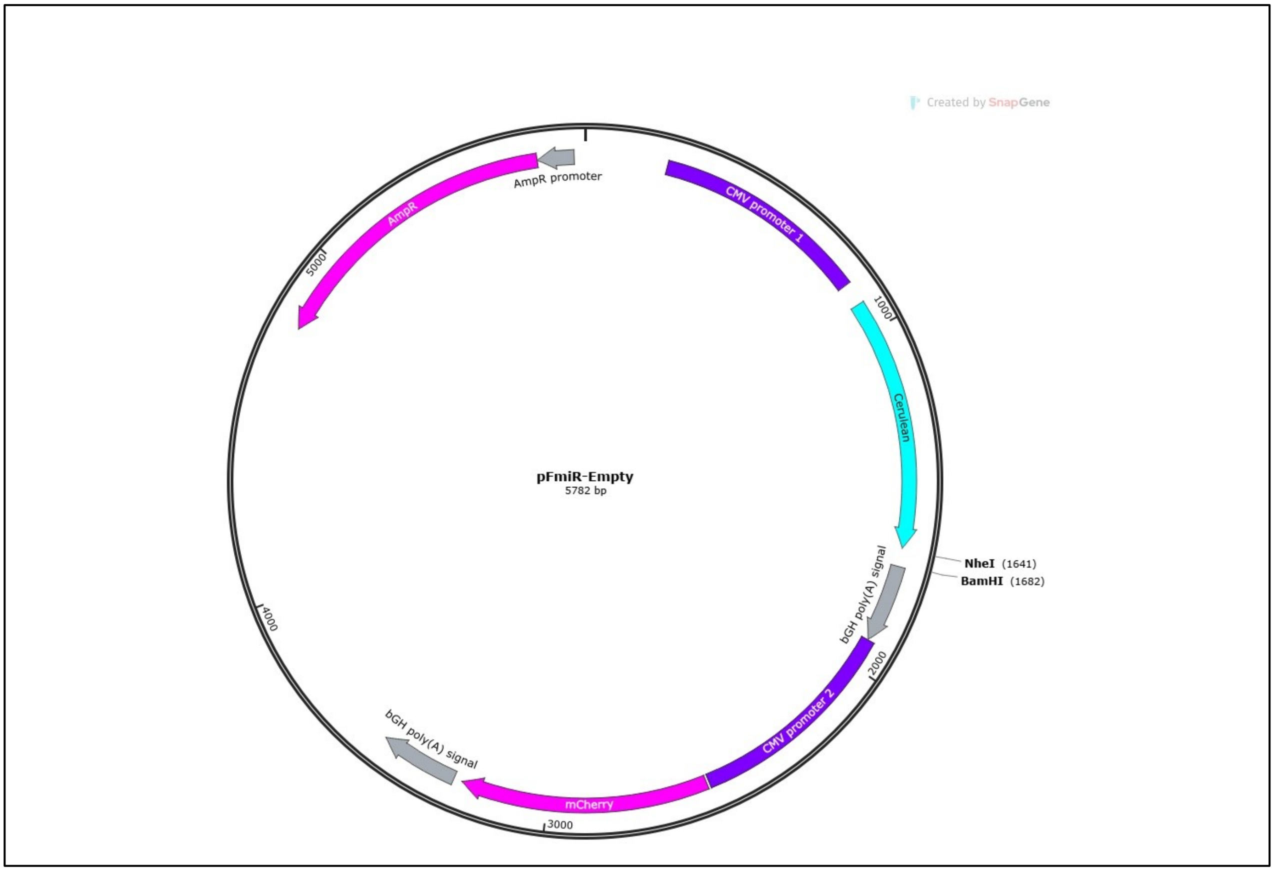

**Figure S1. pFmiR-OGT map** pFmiR-OGT plasmid map. (Created using SnapGene)

5’CACATGATTAAGCCTGTTGAAGTCACTGAGTCAGCATAAATAAAGACTGCACAGGAGAATTACCCCTATACCTGAGCCTCAACCTTCTGGGGGAAAGGGAACTAGATAACATACTTCTTA
CTTGTCTGTACAGTACCTTGTTGCAGATGGGTGATATATAATGGTAATAGAATAGCACAGCCAGACTTGCTTCCTGCATGGTAGGGAGAGACACAAAAGATGGGAAACTGCTTTTCCACAA
GGAATCTCCGTAGAATTTTGCGGCGACCAGATGGTGCATAGGTCTGGAAGGTCTGATCTCCCTTGGTCTTCCATGGGATGGTTAGTGTGGAGGGGAGATATAGATTGTCCGGCCGCTTTG
TGATTCCATGGATTGATTCAGTCTTCTGGATTTTTTTTTCTTTATATTTTGGGTACTGGAGCTTTTAAAAATGTTTGGTTTCAGGTATTTTTATTCATGTGAAGTGTATATGATTCTCTTGAGAT
AAGGTTTTAAGCTAAAATGTTACTCCCTGTTTTAGTTTCTGAACTCTGACAGATTGACAGGGACTTTGCTGGTGTAGTCTTTTTATAGGTTTTATAAACCACTTGAGCCTATATCAGTCGTTTT
AGTGTCTGACCTAATATTTGGAGCTATCAGTGCTTTGTTGATTTAGATGATGACTCAAGATTTTTTCTGGTCCATTTCCCATTTCCTTTTCTTCCCTGACCCCCATACCCTCACCCTTAAAATTCT
CCTGTAACTCAACTAACAAAATCAAGCCTGATTCAAAACATCCTAGGGTGTTTTAAACACACCATCTGGTGCCAAATGAAGATTTTTAGGAGTGATTACTAATTATCAAGGGCACAGTTGTG
GTACTGTCATTGATAATAATATAGTTTTTTTTTTTTTCCTAATTTTGACCTGTTTCACCAGTGTTTTACCCTTGACTGCCCCTTCTATGCTGCTTCCAAAAGTGATAGTGTGTGTAAGATTTTTAC
CTTCCTTTCTAAAGTTTTTTTTTTTTTTTTTTAAGTGAGTCCTGTTCTTCCTATTTCTTTCAGCAGAAATGAAATCCCAGGTAAGTATAAGTATTCAAGTATTTGATCAGTAAGTCACAGTTATCT
CCAGTGCATTAAATAACCTTCATCAAGAAATAGGTTATAGGTAAAATCTCTGAAGGATCATCTATGTATTCAAGTAATTATTTTTTAGATAATAACTGTCTTCTGGACTTGGTCTTGAAGTCT
GTACAGATTCAGCCTCAGTAGTAGCGAACTGCACTGCTGTTTGGTTTGGAGTACAAATTAGACTTATAGTCCTCCTGGAACTTGAGTTATTAAAATCATAGGAATAAAATTATGGGATCTCA
ACAAAGGGTCGAGGGTTTGAGGCTTAAACAAGCCAACATATGAATATATGTTTTGTCTCGCTATACTGCACTTACGCTATCCAGTTGCAGGTAATTTTTTGTCTGCTAGTAGTGTTCTAGATT
ATGTCTTTCCAAAGCGCTGAGGCTGTGCACCTATTCTGTAGTTGCAGCTGATGCCTGAATGTATCCTAGCTGACAAATTATTGATTAATAAGAACTTGAATTTCTGGAAGATTCTTACTGTTA
ACCAAATTTTGAGCAAGGAGTCTCAAAGGTAATTCTGAACCAGAATTACATGTTAATGAACAGTGTACCTTTTAACAGTGTAAATCACGGAATATCCGTGAAGGGATTTCTTAATTTATTTTT
TACCGGTTGATTGAAATATCAGTTAAAGGTTGCCAGCATGGTTGCAGATAAACTGATGTTTGAAATTCGCTGAAATACTTAATGTGGAATAGGATAATATACTTCCAATGCCCTCAAGGCTG
TGACCTTACAGCCATTTTACATAGCACATCATTCCTCCTATAGGGATGAACTTTTTCCTGGCACGAAAAGTAGCCGCTCTGGTTGAAGCTTTGCTTATTGTAACAGGCTTTTATTTCCAGGTA
ATATGTCTTGGAAGACTTAATTCTGATTAGAGATATAGATATTACTGGAAACTAATTGTTTTTTTTCTATTGTACTCTGCTTTATCAAAGAAGTAAAACATTTAAATCGTACTACAGAAATTAA
GATGTTGTCTTGCGATCCTTAATAAATGAATGATTTCCCTTTAATACGGGGATC3’

**Figure S2. OGT 3’UTR sequence.** Sequence contains 2147 base pairs.

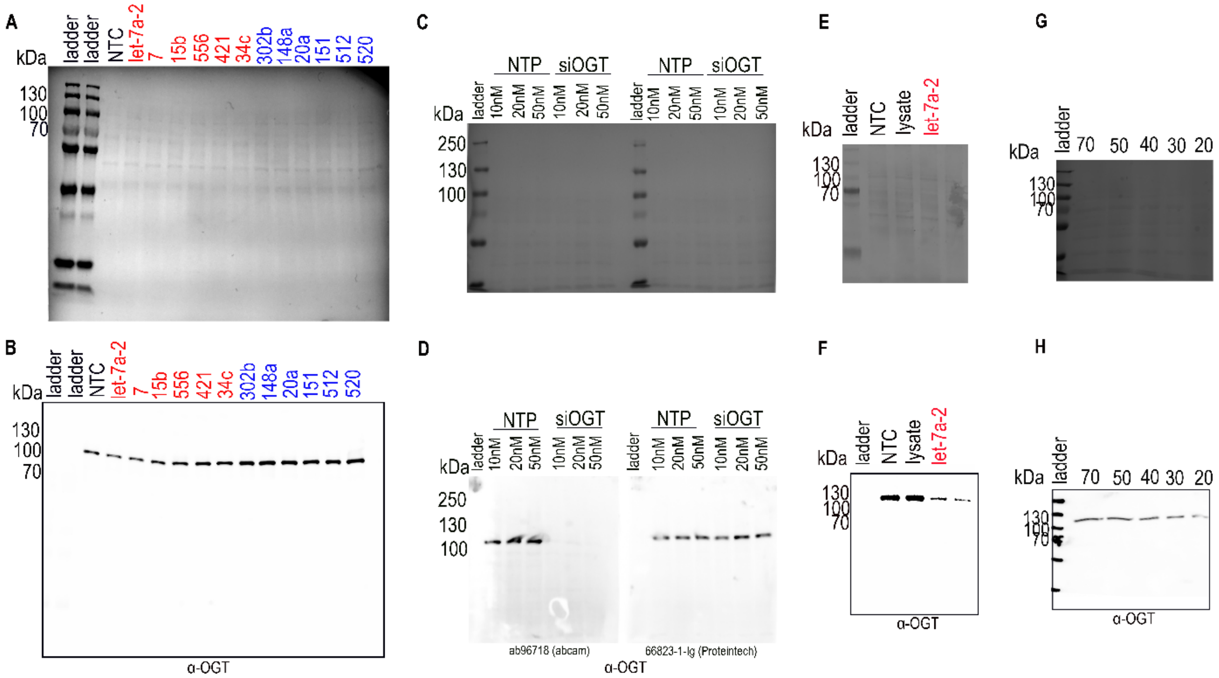

**Figure S3. Ponceau and whole Western blots for data shown in Fig. 3**. (A) Ponceau staining of blot used in **Fig. 3A**. (B) Whole Western blot for data shown in **Fig. 3A** (A549). (C, D) Validation of α-OGT antibodies from Abcam (ab96718) and Proteintech (66823-1). Pooled siRNA against OGT was transfected into A549 using standard protocols. Lysates were run in duplicate on SDS-PAGE and transferred to Nitrocellulose. The blot was then divided in two and stained with α-OGT antibody from Abcam or Proteintech as indicated. Ponceau staining (C) and Western blot (D) showed that while both antibodies detect the expected ~117 kDa OGT band, the band detected by Proteintech could not be silenced by pooled siRNA against OGT. (E, F) Ponceau and Western blot respectively of lysates from A549 cells transfected with 50 nM NTC, let-7a-2-3p (let-7a) or untransfected lysate, showing NTC has no impact on OGT levels. (G, H) Titration of A549 lysate for OGT Western blot analysis. Lysate from A549 (70, 50, 40, 30 and 20µg) was analyzed by Western blot for α-OGT using the Abcam antibody. Ponceau staining (G) and Western blot (H) are shown. We used 20 µg as the optimal concentration for our experiments.

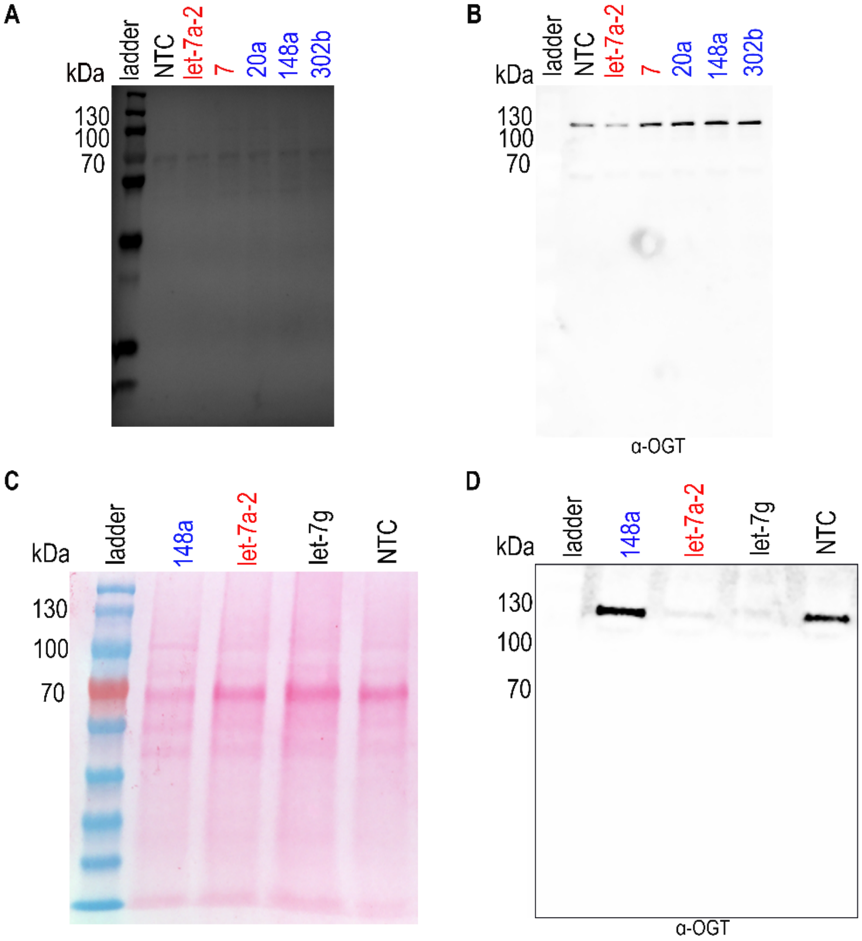

**Figure S4A-D. Ponceau and whole Western blots for data shown in Fig. 4.** (A, C) Ponceau staining of blots shown in **Figs. 4A** (Caco-2, A) **& 4C** (SH-SY5Y, C). (B, D) Whole Western blots for data shown in **Figs**. 4A & C.

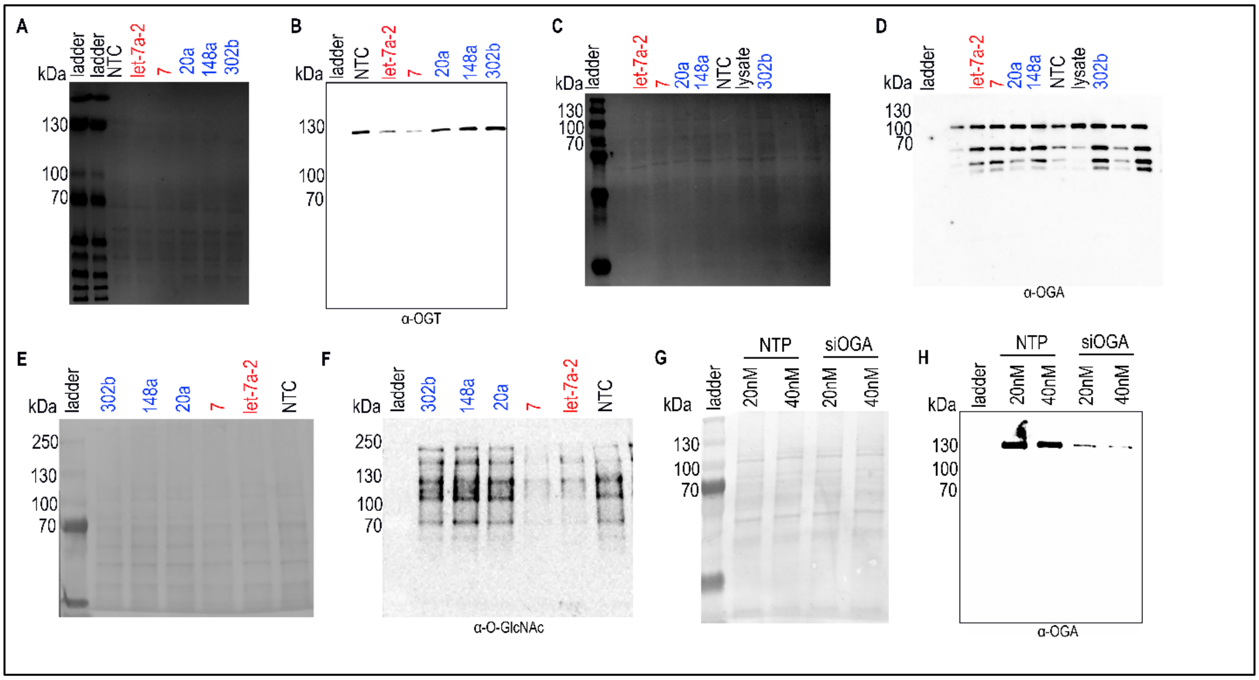

**Figure S5. Ponceau and whole Western blots for data shown in Fig. 5 (A549).** (A, C, E) Ponceau staining of blots used in **Fig. 5A** (OGT), C (OGA) and E (O-GlcNAc). (B, D, F) Whole Western blots for data shown in **Fig. 5A** (OGT), C (OGA) and E (O-GlcNAc). (G-H) Validation of α-OGA antibody (Thermo-Fisher cat # 61425). Pooled siRNA against OGA was transfected into A549 using standard protocols. Lysates were run on SDS-PAGE and transferred to Nitrocellulose and stained with α-OGA antibody. Ponceau staining (G) and whole Western blot (H) confirm the ~115 kDa band as OGA.

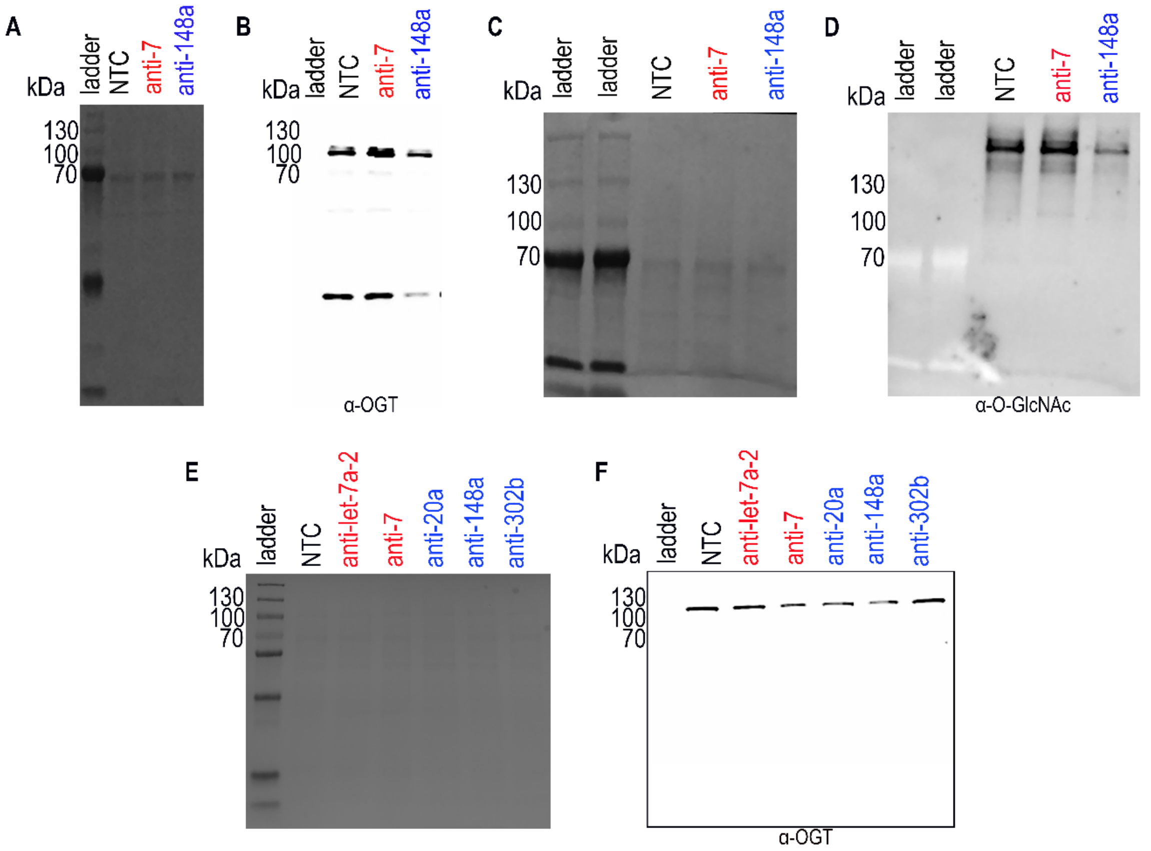

**Figure S6. Ponceau and whole Western blots for data shown in Fig. 6**. (A, C, E) Ponceau staining of blots used in **Fig. 6**, A549 (A, OGT), (C, O-GlcNAc) and Caco-2 (E, OGT). (B, D, F) Whole Western blot for data shown in **Fig. 6**, A549 (A, OGT), (C, O-GlcNAc) and Caco-2 (E, OGT).

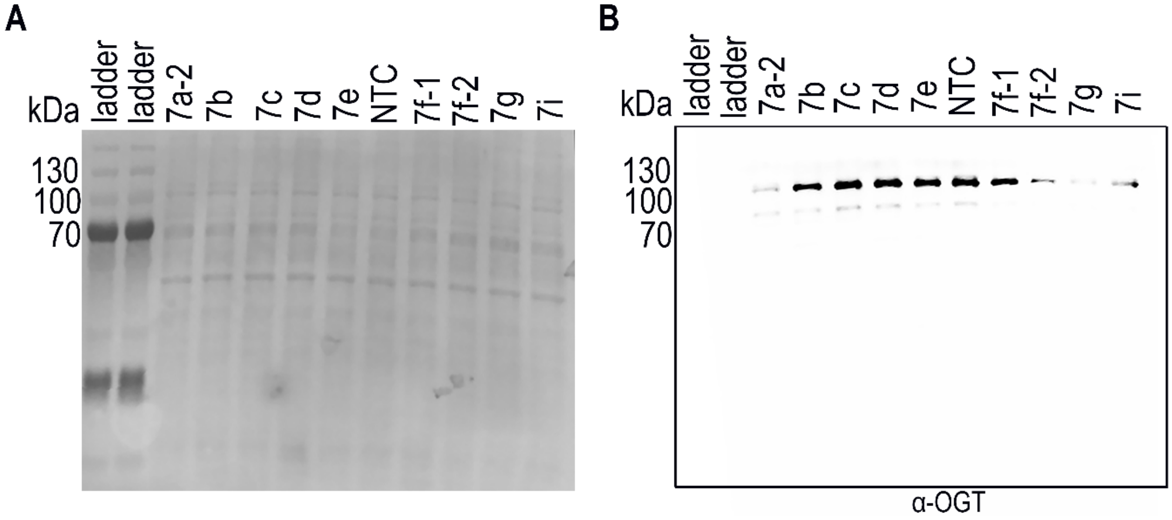

**Figure S7. Ponceau and whole Western blots for data shown in Fig. 7.** (A) Ponceau staining of blot used in **Fig. 7**. (B) Whole Western blot for data shown in **Fig. 7**.

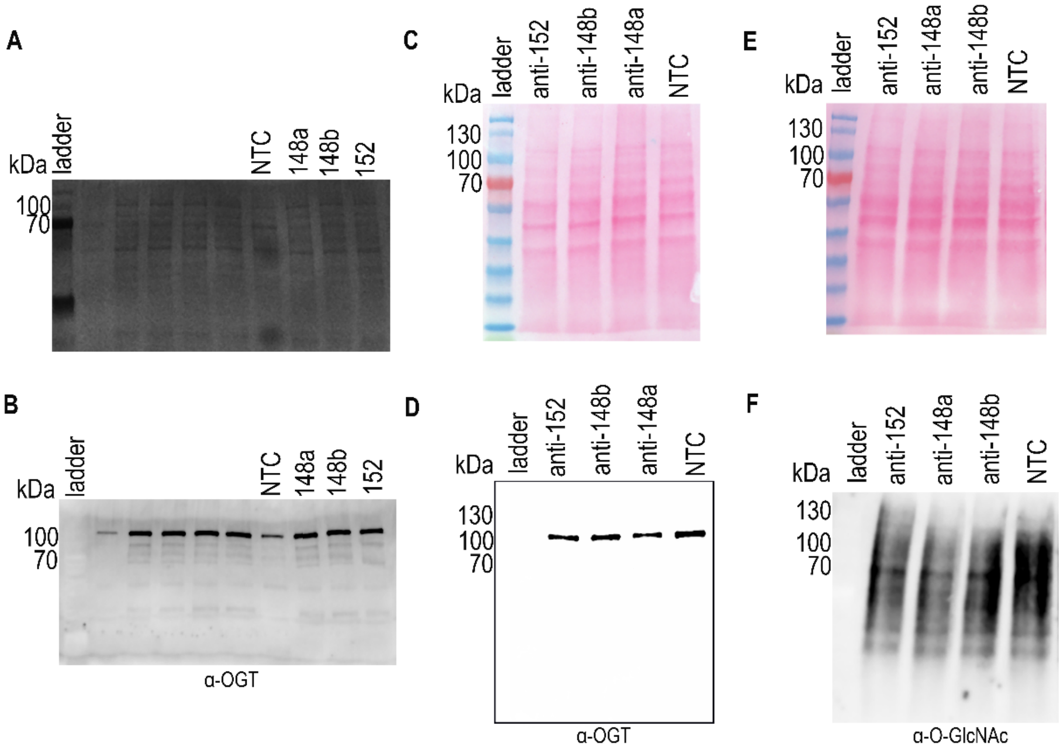

**Figure S8. Ponceau and whole Western blots for data shown in Fig. 9.** (A, C, E) Ponceau staining of blots used in **Fig. 9B** (OGT), **D** (OGT) and **F** (O-GlcNAc). (B, D, F) Whole Western blot for data shown in **Fig. 9B** (OGT), **D** (OGT) and **F** (O-GlcNAc).

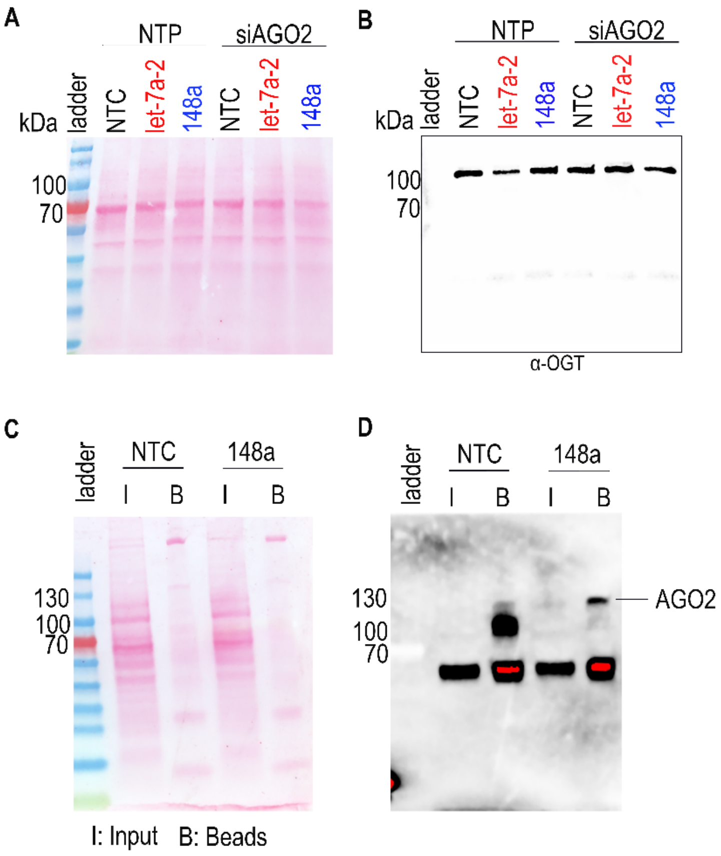

**Figure S9. Ponceau and whole Western blots for data shown in Fig. 10.** (A, B) Ponceau (A) and whole Western blot (B) corresponding to **Fig. 10A**. (C, D) Expanded Ponceau (C) and Western Blot (D) corresponding to **Fig. 10C**. Specificity of the AGO2 antibody was confirmed using AGO2 knockout cell line (not shown).

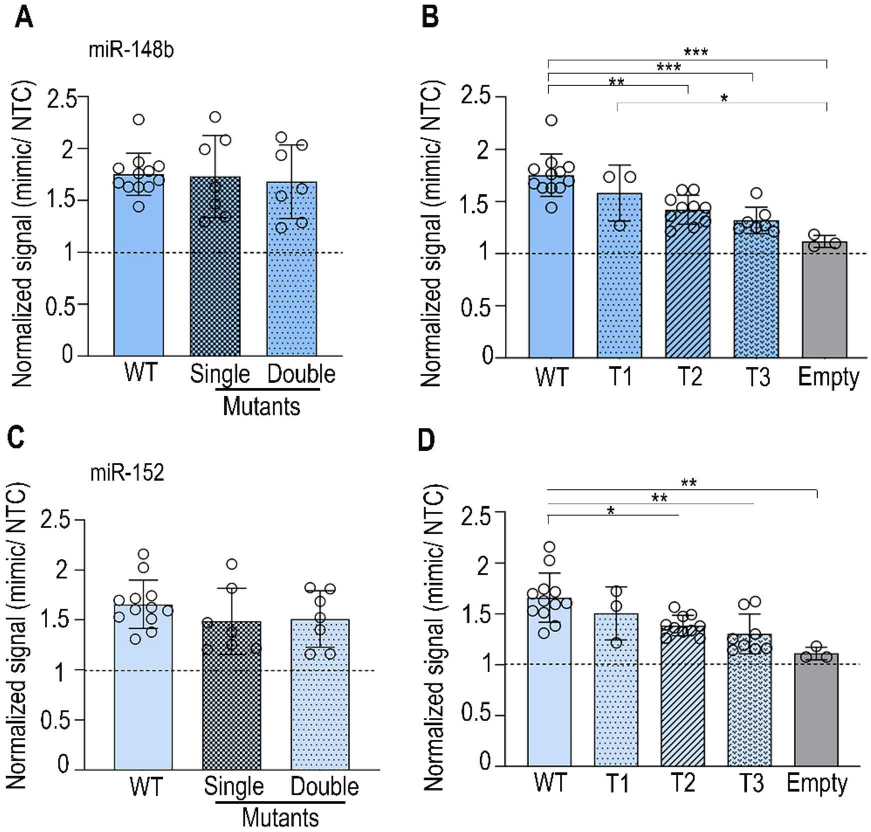

**Figure S10. miR-148b-3p & -152 upregulates OGT through multiple binding sites.** (A) Bar graph shows average data for HEK-293T cells transfected with miR-148b-3p (with WT (wildtype) or MUT (Single, Double) miRFluR sensors. (B) Bar graph shows average data for HEK-293T cells transfected with miR-148b-3p (with WT (wildtype), truncated (T1, T2, T3) or empty pFmiR) miRFluR sensors. (C) Bar graph shows average data for HEK-293T cells transfected with miR-152 (with WT (wildtype) or MUT (Single, Double) miRFluR sensors. (D) Bar graph shows average data for HEK-293T cells transfected with miR-152 (with WT (wildtype), truncated (T1, T2, T3) or empty pFmiR) miRFluR sensors. For all graphs, data was normalized over NTC in that sensor (n≥3 biological replicates, individual replicates indicated). Statistics were done using two-way ANOVA with Tukey's comparison. *p*-values are reported in **Table S2**. *$*p < 0.05$, ** < 0.01, *** <0.001*.

**Table S1. miRFluR assay hits for OGT.** miRNA hits in 90% confidence interval (CI) are represented with down-miRs (red) and up-miRs (blue). Data is normalized to non-targeting control 1 (NTC1). Standard deviation is represented for each miRNA. Complete dataset pre- and post-quality control is shown in **Dataset S1**.

| miRNA name | Normalized to NTC1 | Standard Deviation (SD) |
|---|---|---|
| hsa-let-7a-2-3p | 0.407848803 | 0.221682062 |
| hsa-miR-556-3p | 0.440455857 | 0.055972666 |
| hsa-miR-6757-3p | 0.465658823 | 0.057872106 |
| hsa-miR-4712-5p | 0.487423602 | 0.344660534 |
| hsa-miR-15b-3p | 0.489258739 | 0.345958172 |
| hsa-miR-502-3p | 0.52818451 | 0.373482849 |
| hsa-miR-6736-3p | 0.531461849 | 0.375800277 |
| hsa-miR-421 | 0.534105495 | 0.041326267 |
| hsa-miR-544a | 0.54607173 | 0.386131023 |
| hsa-miR-9500 | 0.54718504 | 0.16181968 |
| hsa-miR-1275 | 0.561759777 | 0.161834604 |
| hsa-miR-6499-5p | 0.566904151 | 0.40086177 |
| hsa-miR-552-3p | 0.568261227 | 0.401821367 |
| hsa-miR-7-5p | 0.571237575 | 0.1705681 |
| hsa-miR-466 | 0.574282062 | 0.272139134 |
| hsa-miR-6854-5p | 0.576104893 | 0.169674841 |
| hsa-miR-4696 | 0.576192475 | 0.407429607 |
| hsa-miR-205-3p | 0.57865791 | 0.043951361 |
| hsa-miR-4476 | 0.586615922 | 0.157833776 |
| hsa-miR-6835-3p | 0.587941417 | 1.25708E-05 |
| hsa-miR-6509-3p | 0.595233326 | 0.004153969 |
| hsa-miR-517c-3p | 0.596323472 | 0.421664371 |

| | | |
|---|---|---|
| hsa-miR-770-5p | 0.602822368 | 0.426259784 |
| hsa-miR-2114-5p | 0.606969223 | 0.318929789 |
| hsa-miR-7705 | 0.609229359 | 0.430790211 |
| hsa-miR-3202 | 0.609418775 | 0.020075816 |
| hsa-miR-627-5p | 0.611593509 | 0.432461917 |
| hsa-miR-501-3p | 0.612656779 | 0.433213763 |
| hsa-miR-624-5p | 0.615083202 | 0.434929503 |
| hsa-miR-4661-3p | 0.621093342 | 0.183865151 |
| hsa-miR-93-3p | 0.622790341 | 0.21250105 |
| hsa-miR-7851-3p | 0.624117778 | 0.441317913 |
| hsa-miR-4711-3p | 0.624140432 | 0.441333932 |
| hsa-miR-1264 | 0.627656088 | 0.007458516 |
| hsa-miR-5579-3p | 0.629592011 | 0.101889379 |
| hsa-miR-362-3p | 0.629652977 | 0.322674846 |
| hsa-miR-628-5p | 0.63603951 | 0.449747851 |
| hsa-miR-320c | 0.639297441 | 0.255422495 |
| hsa-miR-141-5p | 0.640289451 | 0.250557583 |
| hsa-miR-16-5p | 0.642247758 | 0.29779497 |
| hsa-miR-4704-5p | 0.644201265 | 0.105851625 |
| hsa-miR-218-5p | 0.644375384 | 0.455642203 |
| hsa-miR-24-3p | 0.647999787 | 0.144393595 |
| hsa-miR-6505-3p | 0.650908278 | 0.460261657 |
| hsa-miR-6820-3p | 0.651620608 | 0.460765351 |
| hsa-miR-6505-5p | 0.652001327 | 0.046705303 |
| hsa-miR-3164 | 0.663621204 | 0.028610133 |
| hsa-miR-544b | 0.666534654 | 0.125691886 |
| hsa-miR-3118 | 0.667059399 | 0.346027818 |

| | | |
|---|---|---|
| hsa-miR-3941 | 0.668180531 | 0.472474985 |
| hsa-miR-29a-3p | 0.669236685 | 0.473221798 |
| hsa-miR-4729 | 0.674849672 | 0.168111434 |
| hsa-miR-1290 | 0.677044975 | 0.216842478 |
| hsa-miR-664a-3p | 0.679731187 | 0.480642532 |
| hsa-miR-376c-3p | 0.68102029 | 0.481554066 |
| hsa-miR-5190 | 0.681310556 | 0.481759314 |
| hsa-miR-555 | 0.682012106 | 0.119146791 |
| hsa-miR-34c-3p | 0.692901003 | 0.111688529 |
| hsa-miR-891a-3p | 0.693624938 | 0.128351639 |
| hsa-miR-431-5p | 1.637086736 | 1.900670751 |
| hsa-miR-3177-3p | 1.641276516 | 1.916477281 |
| hsa-miR-219b-3p | 1.647543856 | 1.940121696 |
| hsa-miR-3678-5p | 1.648699068 | 1.944479894 |
| hsa-miR-1973 | 1.652530354 | 1.958933955 |
| hsa-miR-4316 | 1.656434153 | 1.973661584 |
| hsa-miR-520f-3p | 1.657964331 | 1.979434392 |
| hsa-miR-376a-3p | 1.661217027 | 1.991705644 |
| hsa-miR-3668 | 1.664364135 | 2.003578548 |
| hsa-miR-518f-5p | 1.666122019 | 2.01021041 |
| hsa-miR-4691-3p | 1.668513822 | 2.019233822 |
| hsa-miR-4707-5p | 1.669032311 | 2.021189895 |
| hsa-miR-5189-5p | 1.671559129 | 2.030722666 |
| hsa-miR-6858-5p | 1.675132836 | 2.044204974 |
| hsa-miR-106a-5p | 1.678439869 | 2.05668122 |
| hsa-miR-519d-3p | 1.684903319 | 2.081065489 |
| hsa-miR-30c-5p | 1.68640455 | 2.086729091 |

| | | |
|---|---|---|
| hsa-miR-4273 | 1.691756503 | 2.106920081 |
| hsa-miR-1539 | 1.69850996 | 2.132398442 |
| hsa-miR-1827 | 1.702934125 | 2.14908922 |
| hsa-miR-6783-3p | 1.703074037 | 2.149617059 |
| hsa-miR-409-3p | 1.705585051 | 2.159090211 |
| hsa-miR-148b-3p | 1.707844434 | 2.167614046 |
| hsa-miR-506-5p | 1.710045123 | 2.175916455 |
| hsa-miR-6790-5p | 1.710474996 | 2.177538212 |
| hsa-miR-3675-5p | 1.716531889 | 2.200388686 |
| hsa-miR-1277-3p | 1.71703799 | 2.20229802 |
| hsa-miR-885-3p | 1.721467701 | 2.219009726 |
| hsa-miR-5700 | 1.724645282 | 2.230997594 |
| hsa-miR-517-5p | 1.725512759 | 2.234270271 |
| hsa-miR-34b-5p | 1.72602242 | 2.236193041 |
| hsa-miR-1281 | 1.727974061 | 2.243555877 |
| hsa-miR-425-3p | 1.732559083 | 2.260853514 |
| hsa-miR-504-3p | 1.734364804 | 2.267665849 |
| hsa-miR-4742-3p | 1.739476725 | 2.286951284 |
| hsa-miR-4784 | 1.741344972 | 2.293999505 |
| hsa-miR-1183 | 1.745163673 | 2.308406089 |
| hsa-miR-221-5p | 1.745553225 | 2.309875728 |
| hsa-miR-3184-5p | 1.746445217 | 2.313240893 |
| hsa-miR-1204 | 1.746759327 | 2.314425918 |
| hsa-miR-2054 | 1.748106675 | 2.319508977 |
| hsa-miR-3180-5p | 1.767257737 | 2.391759031 |
| hsa-miR-1207-5p | 1.769493723 | 2.400194602 |
| hsa-miR-513a-5p | 1.770902333 | 2.405508778 |

| | | |
|---|---|---|
| hsa-miR-3651 | 1.776704528 | 2.427398372 |
| hsa-miR-489-5p | 1.793162869 | 2.489489759 |
| hsa-miR-3677-3p | 1.795055329 | 2.49662933 |
| hsa-miR-1914-3p | 1.797879925 | 2.507285513 |
| hsa-miR-1305 | 1.799625477 | 2.513870852 |
| hsa-miR-527 | 1.804621326 | 2.53271839 |
| hsa-miR-3614-5p | 1.806354105 | 2.539255541 |
| hsa-miR-938 | 1.807219051 | 2.542518669 |
| hsa-miR-520c-3p | 1.808529691 | 2.547463241 |
| hsa-miR-1249-3p | 1.813831942 | 2.567466723 |
| hsa-miR-130b-3p | 1.8172513 | 2.580366731 |
| hsa-miR-942-5p | 1.828348927 | 2.622234078 |
| hsa-miR-1255b-5p | 1.834775362 | 2.646478702 |
| hsa-miR-887-3p | 1.84309143 | 2.677852229 |
| hsa-miR-551b-3p | 1.848772356 | 2.699284315 |
| hsa-miR-93-5p | 1.853466499 | 2.716993628 |
| hsa-miR-548s | 1.85899664 | 2.737856856 |
| hsa-miR-152-3p | 1.859953619 | 2.741467193 |
| hsa-miR-4640-5p | 1.864301177 | 2.757868963 |
| hsa-miR-944 | 1.87310361 | 2.79107737 |
| hsa-miR-920 | 1.876270252 | 2.803023967 |
| hsa-miR-1289 | 1.882198267 | 2.825388233 |
| hsa-miR-3163 | 1.892514676 | 2.864308328 |
| hsa-miR-3652 | 1.896104991 | 2.877853294 |
| hsa-miR-182-3p | 1.91955365 | 2.966316632 |
| hsa-miR-877-5p | 1.927263351 | 2.995402558 |

| | | |
|---|---|---|
| hsa-miR-765 | 1.93218835 | 3.013982803 |
| hsa-miR-6845-5p | 1.934913459 | 3.024263657 |
| hsa-miR-450a-5p | 1.953282447 | 3.09356323 |
| hsa-miR-151b | 1.973086106 | 3.168275299 |
| hsa-miR-190a-5p | 1.986605221 | 3.21927805 |
| hsa-miR-1246 | 1.986711215 | 3.219677926 |
| hsa-miR-3669 | 2.015929331 | 3.329907349 |
| hsa-miR-193a-3p | 2.023929903 | 3.360090626 |
| hsa-miR-3662 | 2.068623283 | 3.528702644 |
| hsa-miR-3670 | 2.072298102 | 3.542566409 |
| hsa-miR-1291 | 2.092464927 | 3.618648574 |
| hsa-miR-1228-3p | 2.127139246 | 3.749462286 |
| hsa-miR-1207-3p | 2.143922867 | 3.812780839 |
| hsa-miR-302b-3p | 2.230272418 | 4.138546582 |
| hsa-miR-1248 | 2.255050399 | 4.232024973 |
| hsa-miR-148a-3p | 2.39718814 | 4.768259452 |
| hsa-miR-520d-3p | 2.400856737 | 4.782099747 |
| hsa-miR-212-5p | 2.54626707 | 5.330680538 |
| hsa-miR-3649 | 3.759749833 | 9.908713739 |

**Table S2.** Statistical significance data for all Figures. Western blot experiments display both one-sample *t*-test and paired Student *t*-test. For sensor data, statistical test is indicated.

| miRNA/Anti-miR & Protein tested | One sample *t*-test | Paired *t*-test | Fig. Number | Cell line |
|---|---|---|---|---|
| **Western Blot Analysis** | | | | |
| **Down-miR (OGT)** | | | | |
| let-7a-2-3p | 0.0344 | 0.083 | Fig. 3B | A549 |
| miR-7-5p | 0.0381 | 0.087 | | |
| miR-15b | 0.739 | 0.663 | | |
| miR-556 | 0.492 | 0.754 | | |
| miR-421 | 0.540 | 0.520 | | |
| miR-34c-3p | 0.649 | 0.843 | | |
| **Up-miR (OGT)** | | | | |
| miR-302b-3p | 0.067 | 0.087 | Fig. 3B | A549 |
| miR-148a-3p | 0.078 | 0.051 | | |
| miR-20a-30 | 0.071 | 0.112 | | |
| miR-151b | 0.107 | 0.155 | | |
| miR-512 | 0.108 | 0.172 | | |
| miR-520c-3p | 0.1766 | 0.271 | | |
| **Down-miR (OGT)** | | | | |
| let-7a-2-3p | 0.0029 | 0.0028 | Fig. 4B | Caco-2 |
| miR-7-5p | 0.131 | 0.137 | | |
| **Up-miR (OGT)** | | | | |
| miR-20a-30 | 0.008 | 0.003 | Fig. 4B | Caco-2 |
| miR-148a-3p | 0.0294 | 0.050 | | |
| miR-302b-3p | 0.3225 | 0.438 | | |

| | | | | |
|---|---|---|---|---|
| **Down-miR (OGT)** | | | | |
| let-7a-2-3p | 0.060 | 0.049 | Fig. 4C | SH-SY5Y |
| **Up-miR (OGT)** | | | | |
| miR-148a-3p | 0.074 | 0.119 | Fig. 4C | SH-SY5Y |
| **Down-miR (OGT)** | | | | |
| let-7a-2-3p | 0.031 | 0.0005 | Fig. 5B | A549 |
| miR-7-5p | 0.521 | 0.280 | | |
| **Up-miR (OGT)** | | | | |
| miR-20a-30 | 0.264 | 0.677 | Fig. 5B | A549 |
| miR-148a-3p | 0.159 | 0.761 | | |
| miR-302b-3p | 0.008 | 0.364 | | |
| **Down-miR (OGA)** | | | | |
| let-7a-2-3p | 0.680 | 0.867 | Fig. 5D | A549 |
| miR-7-5p | 0.121 | 0.200 | | |
| **Up-miR (OGA)** | | | | |
| miR-20a-30 | 0.649 | 0.849 | Fig. 5D | A549 |
| miR-148a-3p | 0.51 | 0.469 | | |
| miR-302b-3p | 0.247 | 0.240 | | |
| **Down-miR (O-GlcNAc)** | | | | |
| let-7a-2-3p | 0.006 | 0.105 | Fig. 5F | A549 |
| miR-7-5p | 0.107 | 0.302 | | |
| **Up-miR (O-GlcNAc)** | | | | |
| miR-20a-30 | 0.061 | 0.115 | Fig. 5F | A549 |
| miR-148a-3p | 0.024 | 0.075 | | |
| miR-302b-3p | 0.070 | 0.009 | | |

| | | | | |
|---|---|---|---|---|
| **Down-miR (OGT)** | | | | |
| Anti- miR-7-5p | 0.578 | 0.665 | Fig. 6B | A549 |
| **Up-miR (OGT)** | | | | |
| Anti-miR-148a-3p | 0.077 | 0.080 | Fig. 6B | A549 |
| **Down-miR (O-GlcNAc)** | | | | |
| Anti- miR-7-5p | 0.751 | 0.812 | Fig. 6D | A549 |
| **Up-miR (O-GlcNAc)** | | | | |
| Anti- miR-148a-3p | 0.019 | 0.023 | Fig. 6D | A549 |
| **Down-miR (OGT)** | | | | |
| Anti-let-7a | 0.958 | 0.974 | Fig. 6F | Caco-2 |
| Anti-miR-7-5p | 0.433 | 0.394 | | |
| **Up-miR (OGT)** | | | | |
| Anti-miR-20a | 0.134 | 0.064 | Fig. 6F | Caco-2 |
| Anti-miR-148a | 0.014 | 0.036 | | |
| Anti-miR-302b | 0.136 | 0.165 | | |
| **Down-miR** | | | | |
| let-7a-2-3p | 0.004 | 0.052 | Fig. 7B | A549 |
| let-7g | 0.0013 | 0.092 | | |
| let-7c | 0.155 | 0.452 | | |
| let-7b | 0.050 | 0.188 | | |
| let-7d | 0.385 | 0.875 | | |
| let-7e | 0.049 | 0.248 | | |
| let-7f-1 | 0.053 | 0.204 | | |
| let-7f-2 | 0.019 | 0.131 | | |
| let-7i | 0.023 | 0.108 | | |

| | | | | |
|---|---|---|---|---|
| **Up-miR (OGT)** | | | | |
| miR-148b-3p | 0.016 | 0.33 | Fig. 9C | A549 |
| miR-152-3p | 0.020 | 0.33 | | |
| miR-148a | 0.012 | 0.35 | | |
| **Anti miR (OGT)** | | | | |
| Anti-miR-148a-3p | 0.019 | 0.001 | Fig. 9E | A549 |
| Anti-miR-148b-3p | 0.113 | 0.095 | | |
| Anti-miR-152-3p | 0.0093 | 0.046 | | |
| **Anti miR (O-GlcNAc)** | | | | |
| Anti-miR-148a-3p | 0.0173 | 0.091 | Fig. 9G | A549 |
| Anti-miR-148b-3p | 0.082 | 0.036 | | |
| Anti-miR-152-3p | 0.046 | 0.137 | | |
| **NTP & siAGO2 (OGT)** | | | | |
| NTP: NTC vs. let-7a-2-3p | 0.001 | 0.327 | Fig. 10B | A549 |
| NTP: NTC vs miR-148a-3p | 0.231 | 0.578 | | |
| siAGO2: NTC vs. let-7a-2-3p | 0.868 | 0.622 | | |
| siAGO2: NTC vs miR-148a-3p | 0.221 | 0.93 | | |
| **Sensor data, Paired *t*-test** | | | | |
| let-7a-2-3p | 0.0000010 | | Fig. 7D | HEK-293T |
| let-7g-3p | 0.00026 | | | |
| let-7c-3p | 0.0070 | | | |
| let-7e-3p | 0.2273 | | | |
| let-7f-1-3p | 0.2019 | | | |

| | | | |
|---|---|---|---|
| let-7i-3p | 0.0220 | | |
| let-7a-2-3p-MUT-A | 0.0143 | Fig. 8E | HEK-293T |
| let-7g-3p-MUT-A | 0.0063 | | |
| let-7a-2-3p-MUT-B | 0.2089 | | |
| let-7g-3p-MUT-B | 0.0351 | | |
| let-7a-2-3p-MUT-A+B | 0.0008 | | |
| let-7g-3p-MUT-A+B | 0.0022 | | |
| **Sensor data, 2-way ANOVA** | | | |
| *miR-148a-3p* | | | |
| WT vs Single Mut | 0.388 | Fig. 11C | HEK-293T |
| WT vs Double Mut | 0.526 | | |
| Single Mut vs Double Mut | 0.964 | | |
| WT vs T1 | 0.2432 | Fig. 11D | |
| WT vs T2 | 0.0359 | | |
| WT vs T3 | 0.0074 | | |
| WT vs empty pFmiR | 0.0026 | | |
| T1 vs T2 | 0.999 | | |
| T1 vs T3 | 0.987 | | |
| T1 vs empty pFmiR | 0.329 | | |
| T2 vs T3 | 0.858 | | |
| T2 vs empty pFmiR | 0.161 | | |

| | | | |
|---|---|---|---|
| T3 vs empty pFmiR | 0.460 | | |
| *miR-148b-3p* | | | |
| WT vs Single Mut | 0.839 | Fig. S10A | HEK-293T |
| WT vs Double Mut | 0.560 | | |
| Single Mut vs Double Mut | 0.880 | | |
| WT vs T1 | 0.376 | Fig. S10B | |
| WT vs T2 | 0.0012 | | |
| WT vs T3 | 0.0004 | | |
| WT vs empty pFmiR | 0.0001 | | |
| T1 vs T2 | 0.811 | | |
| T1 vs T3 | 0.357 | | |
| T1 vs empty pFmiR | 0.021 | | |
| T2 vs T3 | 0.764 | | |
| T2 vs empty pFmiR | 0.078 | | |
| T3 vs empty pFmiR | 0.350 | | |
| *miR-152* | | | |
| WT vs Single Mut | 0.094 | Fig. S10C | HEK- |
| WT vs Double Mut | 0.136 | | |
| Single Mut vs Double Mut | 0.972 | | |
| WT vs T1 | 0.479 | | |
| WT vs T2 | 0.028 | | |
| WT vs T3 | 0.0079 | | |

| | | | |
|---|---|---|---|
| WT vs empty pFmiR | 0.0029 | Fig. S10D | 293T |
| T1 vs T2 | 0.995 | | |
| T1 vs T3 | 0.877 | | |
| T1 vs empty pFmiR | 0.180 | | |
| T2 vs T3 | 0.911 | | |
| T2 vs empty pFmiR | 0.199 | | |
| T3 vs empty pFmiR | 0.477 | | |
| **qPCR, One sample *t*-test** | | | |
| let-7a-2-3p vs NTC | 0.0116 | 3C | A549 (*OGT*) |
| miR-148a-3p vs NTC | 0.0829 | | |
| miR-148a-3p vs NTC | 0.0111 | 10D | 293T AGO2 (*OGT*) |

**Table S3-** Primer sequences for PCR amplification of wild-type (WT) 3'UTR (A), site directed mutagenesis of OGT (B)qPCR primers and truncations generated for OGT 3'UTR.

| (A) Primer | Sequence | |
|---|---|---|
| OGT 3'UTR_Fwd | CCACATGATTAAGCCTGTTG | Primers for cloning OGT in pFmiR sensor |
| OGT 3'UTR_Rev | GATCCCCGTATTAAAGGGAAATC | |
| (B) Primer | Sequence (5'→3') | Mutant |
| OGT-148a-TD-FWD | GCACATCCTAGGGTGTTTTAAACACACCATCT | Single Mutant (2nd CLIP site mutated) |
| OGT-148a-TD-REV | GGCCCGACCGGTAAAAAATAAATTAAGAAATC | |
| 148aclip1_FHN-FWD | agtcCTGTTTGGTTTGGAGTAC | Double mutant (Both CLIP sites mutated) |
| 148aclip1_FHN-REV | cgaGTTCGCTACTACTGAGG | |
| **Primer sequence for qPCR** | | |
| Primer | Sequence | Gene |
| Forward | CACCATCCCAGCTCACTT | OGT |
| Reverse | CAGCTTCACAGCTATGTCTTC | |
| Forward | GGTGTGAACCATGAGAAGTATGA | GAPDH |
| Reverse | GAGTCCTTCCACGATACCAAAG | |
| Forward | CAAGTCGGACAGGAGCAGAAA | AGO2 |
| Reverse | GACCTAGCAGTCGCTCTGATCA | |
| Primers for Truncations of OGT 3' UTR | | |
| Forward | GCGTCAGCTAGCCACATGATTAAGCCTGTTGAAG TC | 1500bp |
| Reverse | GCGTCAGGATCCGCGTAAGTGCAGTATAGCG | |
| Forward | GCGTCAGCTAGCCACATGATTAAGCCTGTTGAAG TC | 1Kb |
| Reverse | GCGTCAGGATCCTGATCAAATACTTGAATACTTAT ACTTAC | |

| | | |
|---|---|---|
| Forward | GCGTCAGCTAGCCACATGATTAAGCCTGTTGAAG TC | 500bp |
| Reverse | GCGTCAGGATCCTAGCTTAAAACCTTATCTCAAG AG | |

**Table S4-** Sample data for select species from Targetscan.[1] Binding site (+1757- 1779 bp) for hsa-let-7a-2-3p/-7g in OGT 3' UTR of the specified species. miRNA binding site is shown (red), non- conserved nucleotides are highlighted in blue.

| **Specie name** | **Sequence (5' ____ 3')** | | | | | | | | | | | | |
|---|---|---|---|---|---|---|---|---|---|---|---|---|---|
| Human | U | G | U | C | U | G | U | A | C | A | G | U | |
| Chimp | U | G | U | C | U | G | U | A | C | A | G | U | |
| Gorilla | U | G | U | C | U | G | U | A | C | A | G | U | |
| Macaque | U | G | U | C | U | G | U | A | C | A | G | U | |
| Tree shrew | U | A | U | C | U | G | U | A | C | A | G | U | |
| Mouse | U | A | U | C | U | G | U | G | U | A | G | U | |
| Rat | U | A | U | C | U | G | U | A | U | A | G | U | |
| Squirrel | U | A | U | C | U | G | U | A | C | A | G | U | |
| Ch. Hamster | U | A | U | C | U | G | U | A | C | A | G | U | |
| Guinea pig | U | A | U | C | U | G | U | A | C | A | G | U | |
| Rabbit | U | A | U | C | U | G | U | C | A | G | C | | |
| Pig | U | A | U | C | U | G | U | A | C | A | G | U | |
| Dog | U | A | U | C | U | G | U | A | C | G | G | U | |
| Cat | U | A | C | U | G | U | A | C | A | G | U | | |
| Horse | U | A | C | C | U | G | U | A | C | A | G | U | |
| Elephant | U | A | U | C | U | G | U | A | C | A | G | U | |
| Dolphin | U | A | U | C | U | G | U | A | C | A | A | U | |
| Whale | U | A | U | C | U | G | U | A | C | A | A | U | |
| Hedgehog | U | A | U | C | U | G | U | A | C | A | G | U | |
| Bat | U | A | U | C | U | G | U | A | U | A | G | U | |
| Armadillo | U | A | U | C | U | G | U | A | C | A | G | U | |
| Opossum | U | A | U | C | U | G | C | U | C | A | - | - | |
| Chicken | U | A | A | U | C | U | G | C | C | U | G | G | U |
| Turkey | U | A | A | U | C | U | G | C | C | U | G | G | U |
| Duck | U | A | A | U | C | U | G | C | C | U | G | G | U |
| Pigeon | U | A | A | U | C | U | G | C | C | U | G | G | U |
| Alligator | U | A | A | U | C | U | G | C | C | U | G | G | U |
| Sea turtle | U | A | A | U | C | U | G | C | C | U | G | G | U |
| Lizard | C | U | G | C | C | U | G | G | U | | | | |
| Platypus | - | - | - | - | - | - | - | - | - | - | - | - | - |

**Table S5-** Sample data for select species from Targetscan.[1] Binding site (+2866- 2886 bp) for let-7a-2-3p/-7g in OGT 3' UTR of the specified species. miRNA binding site is shown (red), non-conserved nucleotides are highlighted in blue.

| Specie | Sequence (5' ____ 3') | | | | | | | | | | | | | | | | | | | | | | | |
|---|---|---|---|---|---|---|---|---|---|---|---|---|---|---|---|---|---|---|---|---|---|---|---|---|
| Human | G | G | U | C | U | U | G | A | A | G | U | C | U | G | U | A | C | A | G | A | U | U | C | A | |
| Chimp | G | G | U | C | U | U | G | A | A | G | U | C | U | G | U | A | C | A | G | A | U | U | C | A | |
| Gorilla | G | G | U | C | U | U | G | A | A | G | U | C | U | G | U | A | C | A | G | A | U | U | C | A | |
| Macaque | G | G | U | C | U | U | G | A | A | G | U | C | U | G | U | A | C | A | G | A | U | U | C | A | |
| Tree shrew | - | U | C | U | U | G | A | A | G | U | C | U | G | U | A | C | **G** | G | A | U | U | C | A | | |
| Mouse | **A** | G | **C** | **U** | U | **C** | **A** | A | A | **C** | U | C | - | - | U | A | **U** | **G** | G | A | U | U | **C** | **G** | |
| Rat | **A** | G | U | **U** | **C** | **C** | **A** | A | A | **A** | U | C | - | - | U | A | **U** | **G** | G | A | U | U | C | A | |
| Squirrel | G | G | U | C | U | U | G | A | A | **G** | **U** | **U** | U | G | U | **G** | **U** | **G** | G | A | U | U | C | A | |
| Ch. Hamster | **A** | **A** | U | C | U | **C** | **U** | A | A | G | U | C | - | - | U | A | **U** | **G** | G | A | U | U | C | A | |
| Guinea pig | G | G | U | U | U | U | G | A | A | G | U | C | U | G | U | A | **U** | **U** | G | A | U | U | C | A | |
| Rabbit | G | G | U | C | U | U | G | A | A | G | U | C | U | G | U | A | **U** | **G** | **A** | A | U | U | C | A | |
| Pig | G | G | U | C | U | U | G | A | A | G | U | C | U | G | U | A | **U** | **G** | **A** | A | U | U | C | A | |
| Dog | G | U | C | U | U | G | A | A | G | A | G | U | C | U | G | U | A | **U** | **G** | G | A | - | U | C | A |
| Cat | | - | - | - | - | U | U | G | A | A | G | U | C | U | G | U | A | **C** | **G** | G | A | - | U | C | A |
| Horse | G | G | U | C | U | U | G | A | A | G | U | C | U | **C** | U | **G** | **U** | **G** | G | A | G | G | U | C | A |
| Elephant | **G** | **A** | U | C | U | U | G | A | A | G | U | C | U | G | U | A | **U** | **G** | **A** | A | U | U | C | A | |
| Dolphin | G | G | U | C | U | U | G | A | A | G | U | C | U | **U** | U | A | **U** | **G** | G | A | U | U | C | A | |
| Whale | G | G | U | C | U | U | G | A | A | G | U | C | U | U | **U** | A | U | **G** | G | A | U | U | C | A | |
| Hedgehog | G | G | U | C | U | U | U | C | A | A | G | U | C | U | G | U | **G** | **U** | **G** | G | A | U | U | C | A |
| Bat | G | G | U | C | U | U | G | A | A | G | U | C | U | **C** | U | **G** | **U** | **G** | G | **G** | U | U | C | A | |
| Armadillo | G | G | U | C | U | U | G | A | A | G | C | C | U | G | U | **C** | **U** | G | G | A | U | U | C | A | |
| Opossum | **U** | **C** | **A** | **A** | **G** | U | **A** | **C** | **G** | **U** | **G** | **A** | U | G | U | **C** | C | A | G | A | U | U | **U** | **G** | |
| Chicken | ------------------------------------------------------- |  |  |  |  |  |  |  |  |  |  |  |  |  |  |  |  |  |  |  |  |  |  |  |  |
| Turkey | ------------------------------------------------------- |  |  |  |  |  |  |  |  |  |  |  |  |  |  |  |  |  |  |  |  |  |  |  |  |
| Duck | ------------------------------------------------------- |  |  |  |  |  |  |  |  |  |  |  |  |  |  |  |  |  |  |  |  |  |  |  |  |
| Pigeon | ------------------------------------------------------ |  |  |  |  |  |  |  |  |  |  |  |  |  |  |  |  |  |  |  |  |  |  |  |  |
| Alligator | A | G | U | G | U | C | C | - | - | - | - | - | - | - | - | - | - | - | - | - | C | A | U | C | |
| Sea turtle | --------------------------------------------------------- |  |  |  |  |  |  |  |  |  |  |  |  |  |  |  |  |  |  |  |  |  |  |  |  |
| Lizard | ---------------------------------------------------------- |  |  |  |  |  |  |  |  |  |  |  |  |  |  |  |  |  |  |  |  |  |  |  |  |
| Platypus | ----------------------------------------------------------- |  |  |  |  |  |  |  |  |  |  |  |  |  |  |  |  |  |  |  |  |  |  |  |  |

**Table S6-** Binding sites of miR-148/152 family with AGO2 protein on OGT in HIT-CLIP studies. Acquired from starBase v2.0.[2,3]

| miRNA | Binding site | Log10 (pval) | Binding Type | Cell/Tissue | Accession |
|---|---|---|---|---|---|
| miR-148/152 family | chrX:71575157-71575271 [+][4] | -4 | Non-canonical | HEK-293T (polysome) | SRR15277844 |
| miR-148/152 family | chrX:71575253-71575292 [+][5] | -5 | Non-canonical | Motor-cortex (grey matter) | GSM1259114 |

**Table S7-** Sites 1-2, CLIP sites, Sites 1, 3-7, RNA hybrid[6] binding site predictions for miR-148a in OGT 3'UTR. Respective binding position in OGT 3'UTR with MFE are represented.

<table>
<tr><th>Sites</th><th>Binding sites of miR-148a in OGT 3'UTR</th><th>Binding position (bp)</th><th>MFE (kcal/mol)</th></tr>
<tr><td>1</td><td>target 5' U CCUA U G U 3'<br>GCA UUCUGUAGU GCA CUGA<br>UGU AAGACAUCA CGU GACU<br>miRNA 3' UUC 5'</td><td>1504</td><td>-25.5</td></tr>
<tr><td>2</td><td>Target: 5' CAGUAGUAGCGAACUGCACUGC 3'<br>: ||| | | ||||||||<br>miRNA : 3' UGUUUCAAGACAUCACGUGACU 5'</td><td>1246</td><td>-15.30</td></tr>
<tr><td>3</td><td>target 5' U C UA CC A 3'<br>CA AGU UCU AGUGCAUU<br>GU UCA AGA UCACGUGA<br>miRNA 3' U U CA CU 5'</td><td>1099</td><td>-20.0</td></tr>
<tr><td>4</td><td>target 5' U GCACAG ACU U U 3'<br>CAAGG UUGUGGU G CAUUGA<br>GUUUC GACAUCA C GUGACU<br>miRNA 3' U AA 5'</td><td>842</td><td>-20.2</td></tr>
<tr><td>5</td><td>target 5' G AC C G 3'<br>ACAGGG UUUG UGGUGUA<br>UGUUUC AGAC AUCACGU<br>miRNA 3' A GACU 5'</td><td>542</td><td>-19.3</td></tr>
<tr><td>6</td><td>target 5' U U G GAA GU U 3'<br>GCA AG UCUG G CUGA<br>UGU UC AGAC C GACU<br>miRNA 3' U A AUCA GU 5'</td><td>276</td><td>-19.4</td></tr>
<tr><td>7</td><td>target 5' C UGA G 3'<br>CUGU AGU CACUGA<br>GACA UCA GUGACU<br>miRNA 3' UGUUUCAA C 5'</td><td>13</td><td>-19.2</td></tr>
</table>

**Table S8-** RNA hybrid[6] binding site predictions for miR-148b in OGT 3'UTR. Respective binding position in OGT 3'UTR with MFE are represented.

| Sites | Binding sites of miR-148b in OGT 3'UTR | Binding position (bp) | MFE (kcal/mol) |
|---|---|---|---|
| **1** | target 5' U CCUA AGUUGCAGCU A 3'<br>GCA UUCUGU GAUGC CUGA<br>UGU AAGACA CUACG GACU<br>miRNA 3' UUC U 5' | 1504 | -21.6 |
| **2** | target 5' G U A UAUGUCUUUCCAAA G 3'<br>UAG GUUCU GAU GCGCUGA<br>GUU CAAGA CUA CGUGACU<br>miRNA 3' U U CA 5' | 1464 | -18.9 |
| **3** | target 5' U GCACAG UAC U U 3'<br>CAAGG UUGUGG UG CAUUGA<br>GUUUC GACACU AC GUGACU<br>miRNA 3' U AA 5' | 842 | -20.6 |
| **4** | target 5' G AC C G 3'<br>ACAGGG UUUG UGGUGUA<br>UGUUUC AGAC ACUACGU<br>miRNA 3' A GACU 5' | 542 | -19.0 |
| **5** | target 5' U U G A GU U 3'<br>GCA AG UCUG GA G CUGA<br>UGU UC AGAC CU C GACU<br>miRNA 3' U A A A GU 5' | 276 | -21.6 |
| **6** | target 5' C U A U G 3'<br>CUG UGA G CACUGA<br>GAC ACU C GUGACU<br>miRNA 3' UGUUUCAA A 5' | 13 | -18.7 |

**Table S9-** RNA hybrid[6] binding site predictions for miR-152 in OGT 3'UTR. Respective binding position in OGT 3'UTR with MFE are represented.

| Sites | Binding sites of miR-152 in OGT 3'UTR | Binding position (bp) | MFE (kcal/mol) |
|---|---|---|---|
| 1 | target 5' U AAUA U G AGACUUAAUU U 3'<br> CCAGGU UGUC UG A CUGA<br> GGUUCA ACAG AC U GACU<br>miRNA 3' AG U G 5' | 1962 | -18.5 |
| 2 | target 5' A UA AGU G U 3'<br> CC UUCUGU UGCA CUGA<br> GG AAGACA ACGU GACU<br>miRNA 3' UUC GU 5' | 1507 | -22.9 |
| 3 | target 5' U UAGAUUA UUUCCAAA G 3'<br> GUUC UGUC GCGCUGA<br> CAAG ACAG CGUGACU<br>miRNA 3' GGUU UA 5' | 1468 | -19.9 |
| 4 | target 5' G CAUAUGAAUAUAU UCGCUAUAC U 3'<br> CCAA GUUUUGUC UGCACU<br> GGUU CAAGACAG ACGUGA<br>miRNA 3' U CU 5' | 1387 | -20.4 |
| 5 | target 5' A GG ACA G U 3'<br> UCAAG C GUUGU GUACUG<br> GGUUC G CAGUA CGUGAC<br>miRNA 3' AA A U 5' | 841 | -19.7 |
| 6 | target 5' U C GAA U G 3'<br> UAAG CUGUU G CACUGA<br> GUUC GACAG C GUGACU<br>miRNA 3' G AA UA 5' | 8 | -18.9 |

**Dataset S1. miRFluR data for pFmiR-OGT**. Tab 1. 1st Dataset pre QC. Tab 2. 2nd Dataset pre QC. Tab 3. 1st Dataset after QC. Tab 4. 2nd Dataset after QC. Tab 5. Final list combined. Tab 6. Final list- 90% Confidence interval (CI) *yellow highlighted rows are the miRNAs tested. Confidence intervals are shown. miRNAs found in literature or those for which CLIP sites exist are indicated.

**Dataset S2. ENCORI starbase miRNAs.** Tab 1. DownmiRs common to our hit list and ENCORI starbase v.2 dataset. Tab 2. UpmiRs common to our hit list and ENCORI starbase dataset v.2.
